# Precise kinematic and muscle recording in freely behaving flies enabled by closed-loop tracking and annotation-free pose estimation

**DOI:** 10.64898/2026.03.11.711180

**Authors:** Sibo Wang-Chen, Victor Alfred Stimpfling, Maite Azcorra, Pavan Ramdya

## Abstract

Understanding the neuromuscular basis for behavior requires measuring both kinematic and physiological data at high resolution in unconstrained conditions: a technically challenging goal. Here we present an integrated experimental-computational pipeline for measuring and quantifying body part kinematics and muscle activity in freely behaving *Drosophila melanogaster*. We first present *Spotlight*, a closed-loop videography system that performs real-time tracking to record untethered flies at high resolution (6 µm/pixel) and high frame rate (330 Hz) while also enabling optical recordings of limb muscle activity via a fluorescent calcium reporter. To analyze these massive datasets without manual image annotation, we introduce *PoseForge*, a synthetic-data-driven framework that exploits morphologically accurate biomechanical simulations to generate synthetic data, and contrastive self-supervised learning to infer 3D keypoints and dense body-part segmentation from a single camera view. Using resulting 3D kinematic data, we can replay recorded behaviors in a biomechanical digital twin, NeuroMechFly, to infer forces generated and experienced by the fly’s limbs. Finally, we illustrate the capability of our system to optically record muscle activity. We show how the legs’ long-tendon muscles activate upon mechanical vibration, possibly to activate gripping and to maintain a stable posture. Taken together, this workflow enables scalable, high-resolution measurement and modeling of unconstrained, natural behavior.

## Introduction

Neural circuits are best understood in the behavioral context in which they operate ^1,2^, namely in naturalistic, minimally constrained settings ^3,4,5,6^. Nevertheless, in neuroscience, kinematic measurements are often studied in constrained animals due to the fundamental trade-off between physiological access and behavioral naturalism: with few exceptions ^7,8^, invasive electrophysiological or optical recordings require some tethering or restraint. Thus, an increasingly large gap has formed between the impressively high resolution of connectomes or *structural* maps of neural circuits ^9,10,11,12^, compared with the relatively coarse and unnatural (i.e., constrained) measurements of the *dynamical* behaviors these neural circuits support. This gap is particularly striking in *Drosophila melanogaster*, where recent connectomic datasets ^13,14,15,16,17^ combined with high genetic tractability create exciting opportunities for connectome-constrained modeling ^18,19,20^ of sensorimotor control, yet the detailed behavioral data needed to constrain and validate such models are still lacking.

Here, we report an integrated set of experimental and computational methods that help to reduce this gap. First, we present *Spotlight*, a platform for high-resolution (5–10 µm per pixel) and high frame rate videography of both behavior (300–400 Hz) and muscle activity (30–60 Hz) in freely moving flies (**Video 1, Video 2**). *Spotlight* performs closed-loop camera tracking of the animal, enabling high magnification without restricting the fly’s range of motion or influencing its biomechanics. Unlike approaches with comparable spatiotemporal resolution ^21,22^ that use high-speed cameras that can only record for a few seconds, *Spotlight* processes data streams in real time, allowing long-duration recordings. As muscles constitute the final stage of motor control, measuring their activity provides a direct readout of how neural commands are translated into movement. We therefore extended *Spotlight* to perform simultaneous calcium imaging of muscle activity across multiple limbs. These recordings require no surgical preparation, as muscle fluorescence is imaged directly through the intact cuticle. To our knowledge, this is the first non-invasive recording of limb muscle activity during free behavior.

Analyzing large datasets generated by *Spotlight* and other behavior recording systems requires detailed 3D reconstruction, but existing laboratory animal pose estimation methods typically rely on labor-intensive manual labeling and output sparse keypoint positions ^23,24,25,26^. To overcome these limitations, we developed *PoseForge*, an annotation-free pose estimation framework that eliminates the need for manual image annotation. Instead, we rendered recorded animal poses in NeuroMechFly, an anatomically realistic biomechanical model of *Drosophila* that provides full access to ground-truth kinematic states. We then trained an image-to-image translation model to match the image features of *Spotlight* recordings ^27,28,29^ while preserving simulated poses. This produced synthetic training data paired with precise ground-truth kinematics directly read out from the simulation. To reduce sensitivity to artifacts in our synthetic data, we pretrained an image encoder with contrastive self-supervised learning to extract pose features while ignoring generator-specific noise. Finally, we trained pose estimation models that predict both 3D keypoint positions (from *Spotlight*’s single bottom camera view) and dense, pixel-level segmentation maps of body parts. Together, *PoseForge* enables scalable inference of detailed 3D and dense kinematics.

Detailed kinematics from *Spotlight* and *PoseForge* provide high-quality ground truth for *in silico* studies of animal behavior, which critically rely on accurate, high-resolution data. To demonstrate this, we replayed recorded sequences of joint angles in NeuroMechFly, our fly biomechanical model, and inferred latent dynamical variables including joint torques and ground reaction forces through simulation ^30,18^. Incorporating physics simulation alongside direct experimental measurements connects *kinematics* (observable motion) to inferred *dynamics* (forces generated and experienced by the animal), supporting downstream efforts including categorizing the roles of sensory inputs ^31,32^, relating muscle dynamics to mechanical output ^33,34,35^, modeling neural circuits for sensorimotor control ^19,36,20^, and designing bio-inspired robotic controllers ^37,38,39^.

In parallel, dense body part segmentation maps from *PoseForge* provide regions of interest that can be used to extract fluorescence traces of muscle activity from *Spotlight* recordings. As a demonstration, we quantified how the legs’ long-tendon leg muscles are recruited during mechanical vibration of the arena—a pattern that we noticed only through whole-body-scale behavior and muscle recording in freely behaving animals.

Taken together, these components establish an experimental-computational loop for studying freely moving flies: closed-loop recordings yield high-resolution measurements of unconstrained behavior and muscle activity; annotation-free pose estimation transforms raw recordings into rich kinematic descriptions; *in silico* replay infers underlying forces from movements; and muscle activity measurements help characterize the biomechanical mechanisms supporting limb kinematics. Although our tools were developed for investigating *Drosophila*, the core strategies of our framework—closed-loop imaging for measuring naturalistic behavior, synthetic data for analyzing experimental recordings, dense kinematic categorization, and replay using a digital biomechanical twin—generalize to other organisms and preparations. More broadly, this work illustrates how co-designing experimental systems and computational analysis pipelines can better enable the study of behavior and biomechanics within a comprehensive and quantitative framework.

## Results

### High-resolution, high-frequency recording of freely moving flies

*Drosophila* has long been a powerful model organism because its small size, short duration life cycle, genetic tractability, and, more recently, large-scale connectomics. This same small size, however, makes it technically challenging to record unconstrained fly behavior at high resolution. Recordings of freely behaving flies therefore tend to have either low spatial resolution (e.g., ~ 33–70 pixels per body length ^40,41,42^), a low frame rate (e.g., 1–60 Hz ^42,43^), or short duration (often only seconds) when high-speed cameras cannot process buffered data in real time ^21,22^. Although coarse body postures can be captured at this moderate resolution, this does not enable accurate estimation of joint rotations along the full limb kinematic chain. In fixed camera setups, increased magnification can improve spatial resolution but reduces the observable field of view and arena size. Temporal resolution presents another challenge: flies’ limbs can step at frequencies of up to 20 Hz ^44^, so even 200 Hz recordings can capture as few as 10 frames per step cycle (often only 2–3 frames per leg swing). Higher spatiotemporal resolution can be achieved in tethered preparations (e.g., ~ 500 pixels per body length, up to 180 Hz ^45,46^), but tethering the animal—typically over a spherical treadmill—substantially alters biomechanics and sensory feedback (e.g., gravity loading and proprioception of leg placement). These constraints motivate the development of recording approaches that can combine high spatiotemporal resolution, long duration, with minimal behavioral restriction. Such recordings enable detailed measurements of limb kinematics, multi-joint and multi-leg coordination, and subtle biomechanical effects such as joint compliance and the deformation of soft-segments (e.g., tarsi of the legs), while also supporting the study of long-timescale ethological behaviors like navigation and spontaneous exploration.

To address this need, we developed *Spotlight*, a closed-loop experimental platform that records freely behaving flies at 5–10 µm per pixel (~250–500 pixels per body length) and 300–400 Hz (**Figure 1a,b,e, Video 1 left**). This is accomplished by using a macro lens mounted on a camera that performs closed-loop tracking of the fly in real time (**Video 2**). This design confers several advantages: first, the camera can be placed closer to the animal, improving effective magnification and light collection. Second, the fly remains at a nearly constant distance from the lens, permitting a larger aperture with a shallower depth of field without compromising focus. Third, tracking decouples optical resolution from arena size, enabling recordings in large arenas (up to 150 × 150 mm^2^ in the current design; **Extended Data Fig. 1b**). An inwardly pitched ring light around the lens provides uniform but high-contrast illumination, and the light strobes synchronously with camera exposures to reduce heating. We developed optimized open-source software that enables sustained, high-throughput image acquisition and real-time tracking (**Figure 1c**). The system can be configured for different experimental demands. In the experiments described in this article, data were collected at 5.8 µm pixel size and 330 Hz in a 72 × 48 mm^2^ arena. Optionally, an LED can be installed above the arena for optogenetic activation ^47^ or silencing ^48^ (**Extended Data Fig. 1**).

**Figure 1:**
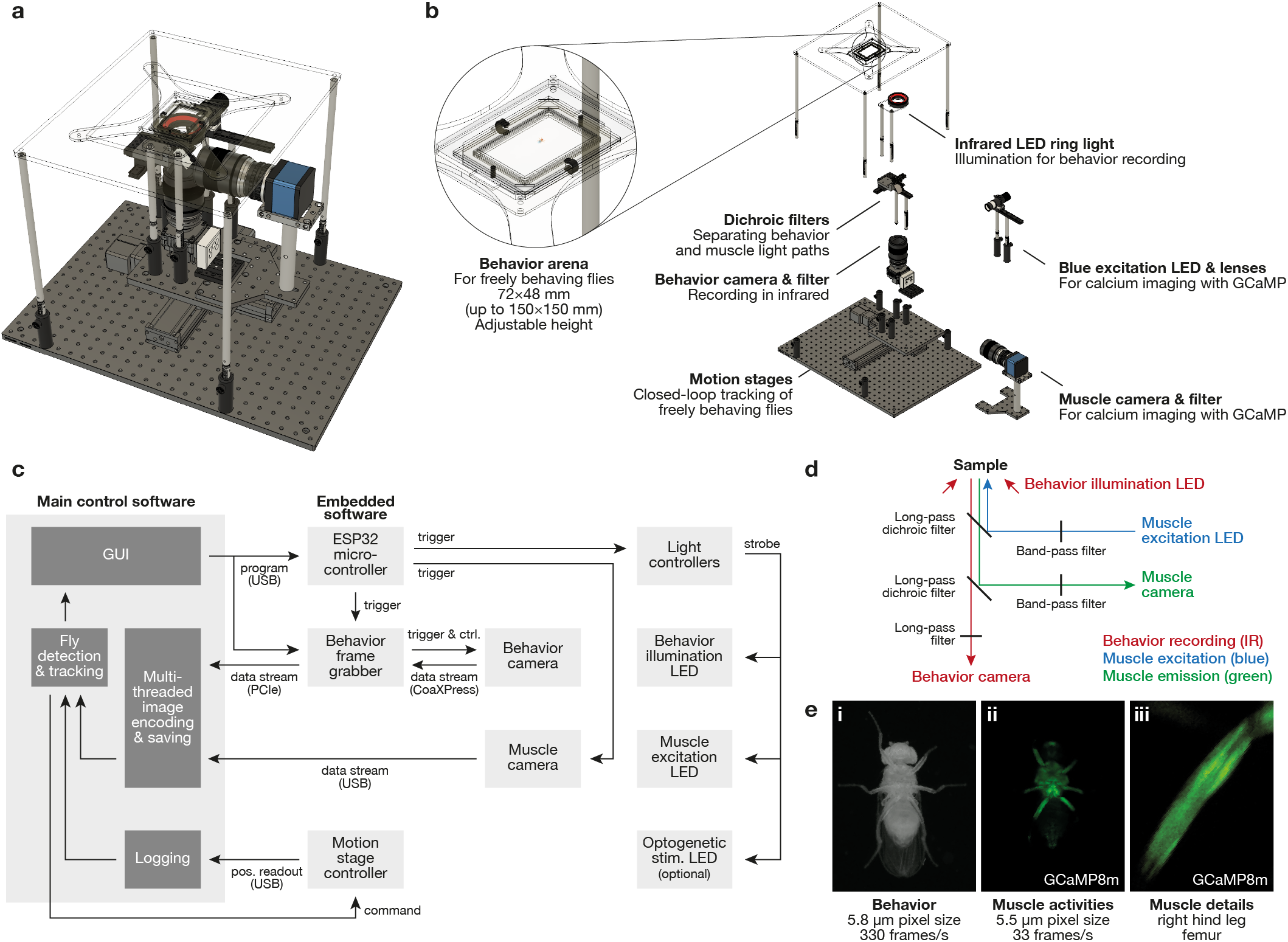
Overview of the *Spotlight* recording system. **(a)** Mechanical design of the experimental system. **(b)** Exploded view of the main optomechanical components, including a zoomed-in view of the arena in which flies can behave freely. **(c)** Simplified block diagram of the software describing the data acquisition, processing, and closed-loop tracking. **(d)** Schematic diagram of optical paths. **(e)** Example recordings from the *Spotlight* system, including (i) a behavior image (recorded using infrared light), (ii) a muscle fluorescence image (recorded using blue excitation and green emission), and (iii) a zoomed-in view of the muscles within a segment of the femur.

### Simultaneous behavior and muscle recording across limbs

Physiological recording of muscles in behaving adult flies is technically very challenging ^33,34,49,50,51^ and, in particular, simultaneous recording across many muscles and limbs in behaving animals has been impossible until now. To connect behavioral *kinematics* (motion) with *dynamics* (force generation and muscle activities), we added a second camera to *Spotlight* to perform calcium imaging of muscles in parallel with behavioral recording (**Figure 1b, Video 1 right**). We expressed the fluorescent calcium indicator GCaMP8m ^52^ in muscles. In this way, increases in intracellular calcium ion concentration associated with muscle activation ^53^ increase muscle fluorescence, which we can measure optically. We recorded behavioral camera frames using infrared illumination, reserving blue excitation and green emission bands for GCaMP8m. Optical paths were separated by spectral bands using dichroic mirrors and filters (**Figure 1d, Extended Data Fig. 1a**). Excitation light was strobed to reduce heating and overstimulation of the fly. The two cameras were hardware-synchronized and spatially registered using custom software (**Figure 1c**). This configuration supports muscle recordings at up to 5.5 µm-per-pixel resolution at ~ 60 Hz; here we used 5.5 µm and 33 Hz. Given the reported GCaMP8m half-decay time (134 ms, or ~ 4.4 frames at 33 Hz, albeit measured in neuronal cultures ^52^), this frame rate balances temporal resolution with reduced stimulation. These recordings resolve leg-segment-level fluorescence and, in some cases, individual muscles within a segment (**Figure 1e**). For the first time, this approach enables non-invasive, simultaneous muscle recordings across all limbs during free behavior.

### Domain transfer between experimental data and simulated images

Deep learning-based pose estimation has revolutionized quantitative behavioral analysis in systems neuroscience ^23,24,25^, but acquiring kinematic descriptions beyond sparse keypoint positions and joint angles remains challenging. This is partly because many quantities of interest are difficult—sometimes impossible—to label reliably by hand (e.g., 3D pose from a single camera view) or prohibitively expensive to annotate at scale (e.g., pixel-wise mapping to body parts or full-body surface correspondence). Synthetic datasets offer an attractive alternative because they can be generated automatically with deterministic ground truth labels. In the broader computer vision field, particularly dense human pose estimation ^54,55^, synthetic data is now widely used to reduce human effort and improve robustness through data randomization ^28,27^. In parallel, anatomically realistic biomechanical models have recently been developed for several animals ^56^, including *Drosophila* ^30,18,19^. Together, there is now an emerging opportunity for annotation-free behavior analysis based on simulation-derived synthetic data paired with kinematic states.

A major barrier, however, is the simulation-to-experiment domain gap: body model renderings still lack the photorealism required to directly mirror real experimental data. To bridge this gap, we trained a generative deep learning model to translate simulated renderings into *Spotlight*-like images while preserving poses (**Figure 2a, Video 3 top & middle**). To accomplish this, we replayed 3D kinematics from a published dataset ^57^ in NeuroMechFly ^30,18^ and rendered images using a virtual camera approximating the *Spotlight* setup. This dataset contains tethered recordings reconstructed from seven camera views, enabling accurate ground truth joint angles estimation ^24^. Notably however, because the fly is tethered on a spherical treadmill, these kinematics cover a biased prior for free locomotion—an important limitation to address in future work.

**Figure 2:**
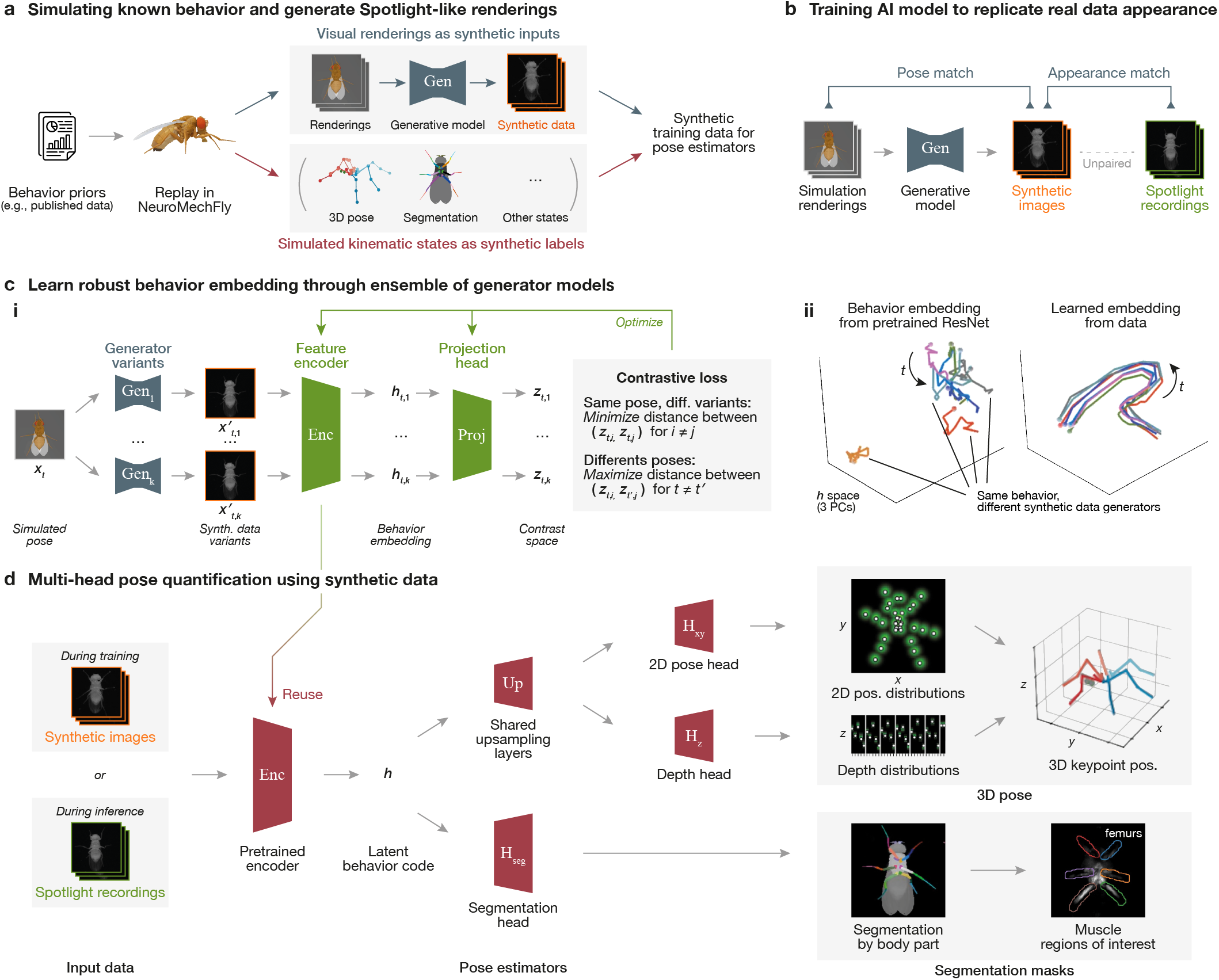
Overview of *PoseForge*, a computational framework that uses synthetic data to perform annotation-free pose estimation. **(a)** Measured behaviors are replayed in a neuromechanical model of adult *Drosophila melanogaster*, NeuroMechFly. This generates renderings with full ground-truth kinematic states (3D keypoints and body part segmentation masks). A generative image-to-image translation model transforms simulation renderings into *Spotlight*-like images, serving as synthetic training inputs while poses read out from the simulation serve as training labels. **(b)** The image generation model Gen is trained to preserve pose from simulated renderings while matching the visual appearance of *Spotlight* data. The training algorithm matches generator output with *Spotlight* recordings only by *distribution*; pairwise correspondence between the two datasets is not required. **(c**,**i)** An image encoder (Enc) is pretrained to extract robust pose representations. For each time point *t* in simulation, multiple versions of the same pose (***x***_*t*,1_, …, ***x***_*t,k*_) are generated using different generator variants (Gen_1_, …, Gen_*k*_). The encoder maps each image to a latent representation ***h***_*t,i*_, which is further mapped to a contrastive embedding ***z***_*t,i*_ by a projection head (Proj). During training, positive pairs (same pose, different generators) are pulled together while negative pairs (different poses) are pushed apart. **(c**,**ii)** Without pretraining (left), generator identity dominates variability in behavior embeddings (in ***h***-space) even for the same behavioral sequence. After pretraining (right), embeddings become largely invariant to generator identity, and instead reflect the smooth evolution of poses through time. See also **Supplementary Video 4. (d)** Pose information is derived through a multi-head decoding architecture. The pretrained encoder extracts a latent code ***h***. This latent code is then upsampled to an intermediate feature map, followed by *x*-*y* and depth heads that together estimate 3D keypoint positions. Concurrently, a segmentation head (H_seg_) upsamples ***h*** into pixel-wise body part masks, thereby defining regions of interest on muscle fluorescence images.

With these NeuroMechFly renderings and real *Spotlight* recordings, we used Contrastive Unpaired Translation (CUT) ^29^ to map simulated images to the experimental domain. CUT combines generative adversarial learning ^58^, which enforces realism in the output domain, with contrastive constraints ^59^ that preserve local structures and reduce hallucinations—a common pitfall for generative AI models. This process produces synthetic *Spotlight*-like images that retain simulated ground truth (**Figure 2b**), enabling scalable training data generation for downstream computer vision tasks. Importantly, this pipeline is not data-limited, as both NeuroMechFly renderings and unannotated *Spotlight* recordings can be generated at low cost and in arbitrary volumes (although resampling of existing kinematic datasets might be necessary to obtain more simulated renderings).

### Self-supervised learning of pose features using synthetic data

Although the synthetic data generator produces visually plausible images, artifacts are inevitable in AI-generated data. Training pose estimators directly on such data can cause models to exploit these artifacts instead of relying on true anatomical features, limiting generalization to real recordings.

To address this issue, we used contrastive self-supervised learning to obtain a latent embedding of pose that encodes body configuration but is insensitive to artifacts and domain-specific noise. We started by training multiple variants of the synthetic data generator (**Figure 2c,i**) with different hyperparameters (e.g., model size, loss weights, training batch size), thereby inducing distinct statistical biases and artifacts in the outputs. We next sought to leverage this diversity across generator variants through ensemble learning, which improves robustness by averaging over per-model biases ^60^. However, because photorealistic images are highly nonlinear at the pixel level, simple averaging of generator outputs is not meaningful. Instead, we trained a convolutional network to encode each image into a feature vector using an InfoNCE ^59^/SimCLR ^61^-style contrastive objective. This objective function pulls images depicting the same underlying pose but produced by different generators (“positive pairs”) together in a latent space, while pushing images depicting different poses (“negative pairs”) apart. As in standard contrastive learning, we applied the objective after a small secondary encoder (“projection head”) to preserve a general-purpose representation that is not overly specialized for the contrastive task.

Unlike conventional contrastive learning methods that define positive and negative pairs through random input transforms (e.g., cropping, resizing, color jittering) ^61^ or human-defined semantic classes ^62^, our positives are defined by shared ground-truth pose across generator variants, conceptually combining domain randomization with input augmentation. This encourages the encoder to emphasize behaviorally relevant geometric features rather than visual idiosyncrasies. As a result, behavior trajectories in the learned latent space are substantially less sensitive to generator identity compared to a generically pretrained ResNet model ^63^. Instead, they vary smoothly over time, dominated by pose changes (**Figure 2c,ii; Video 3 bottom**). We used this contrastively pretrained encoder as the backbone for subsequent pose estimators presented henceforth.

### Training pose estimators without human annotation

Using synthetic images and labels from simulation, and initializing the image encoder with contrastively pretrained weights, we trained pose estimation models for *Spotlight* without human annotation. This enables prediction not only of sparse keypoint positions (e.g., at limb joints), but also richer kinematic descriptors (e.g. dense pixel-level labels) that are impractical to annotate manually. We implemented a *multi-head* architecture in which task-specific *heads* decode a shared latent representation into different output forms (**Figure 2d**). The pretrained encoder that produces these latent codes accounts for ~ 70% of parameters in the pose estimation neural networks, meaning most model capacity is already learned during self-supervised pretraining (**Extended Data Fig. 2**). As a result, pose model training primarily focuses on fitting lightweight task-specific heads, substantially reducing the learning burden.

For 3D keypoint estimation, the encoder output is first upsampled into an intermediate feature map. To preserve spatial detail needed for accurate localization, we used U-Net ^64^-style skip connections that propagate high-resolution features from the downsampling pathway directly to later decoding layers, thereby retaining information that is otherwise lost in the encoding process. A *2D pose head* (H_*xy*_) then predicts per-keypoint probability heatmaps, while a *depth head* (H_z_) predicts per-keypoint depth distributions. The ground truth for depth (i.e., distance from camera) is uniquely gained through simulation in NeuroMechFly, where pose is fully specified subject to biomechanical positions. We combined these outputs to reconstruct 3D keypoint coordinates (**Video 4**). Notably, the 2D pose and depth pathways diverge only after the intermediate feature map and therefore share the vast majority of network parameters (~ 98%; **Extended Data Fig. 2**). This shared representation helps enforce geometric consistency even though 2D and depth coordinates are predicted separately. Although predictions are made over discretized bins, coordinates are obtained using soft-argmax (weighted centroid). Since keypoint likelihood varies smoothly, this effectively provides sub-bin localization precision. To validate depth inference, we computed leg segment lengths and observed that they remain approximately constant over time and left-right symmetrical, as expected (**Extended Data Fig. 3**). This geometric consistency, together with direct validation of *x*-*y* accuracy, supports the reliability of predicted *z* coordinates.

In a separate pathway, a *segmentation head* (H_seg_) decodes the shared embedding into pixel-level body-part masks (**Video 5**) also using U-Net-style skip connections to retain high-frequency spatial information (**Figure 2d**). This dense representation assigns each pixel to a body segment rather than detecting sparse landmarks. In *Spotlight* recordings, these masks enable automated definition of muscle regions of interest for extracting muscle fluorescence signals.

Although all pathways share the same pretrained encoder, encoder weights were fine-tuned during task-specific training at a reduced learning rate (10% of the remaining model), preserving invariances learned through pretraining while still allowing adaptation to specific sub-tasks.

### Inferring unmeasured quantities using replay in a biomechanical digital twin

Although *Spotlight* and *PoseForge* enable high-resolution behavioral quantification, recordings alone do not directly give access to all biomechanical variables. Fundamentally, kinematic tracking captures pose and movement, but not dynamical quantities such as joint torques, ground reaction forces, or mechanical power. These variables are essential for understanding how motor commands *generate* the observed movements.

Biomechanical digital twins provide a surrogate with which one can infer such unmeasured quantities by replaying measured kinematics in a physics simulator ^56^. To illustrate this for freely behaving flies, we first converted tracked 3D keypoint trajectories into joint angles using SeqIKPy ^65^, a sequential inverse kinematics pipeline that fits joint configurations under biomechanical constraints (**Figure 3a**). This step enforces fixed leg segment lengths and restricts joint rotations to anatomically supported degrees of freedom (7 per leg, rather than 12 for unconstrained 3-axis rotations at every joint) (**Figure 3c**). Unlike previous work ^30^, we optimized joint angles to fit all tracked keypoints rather than the claws alone.

**Figure 3:**
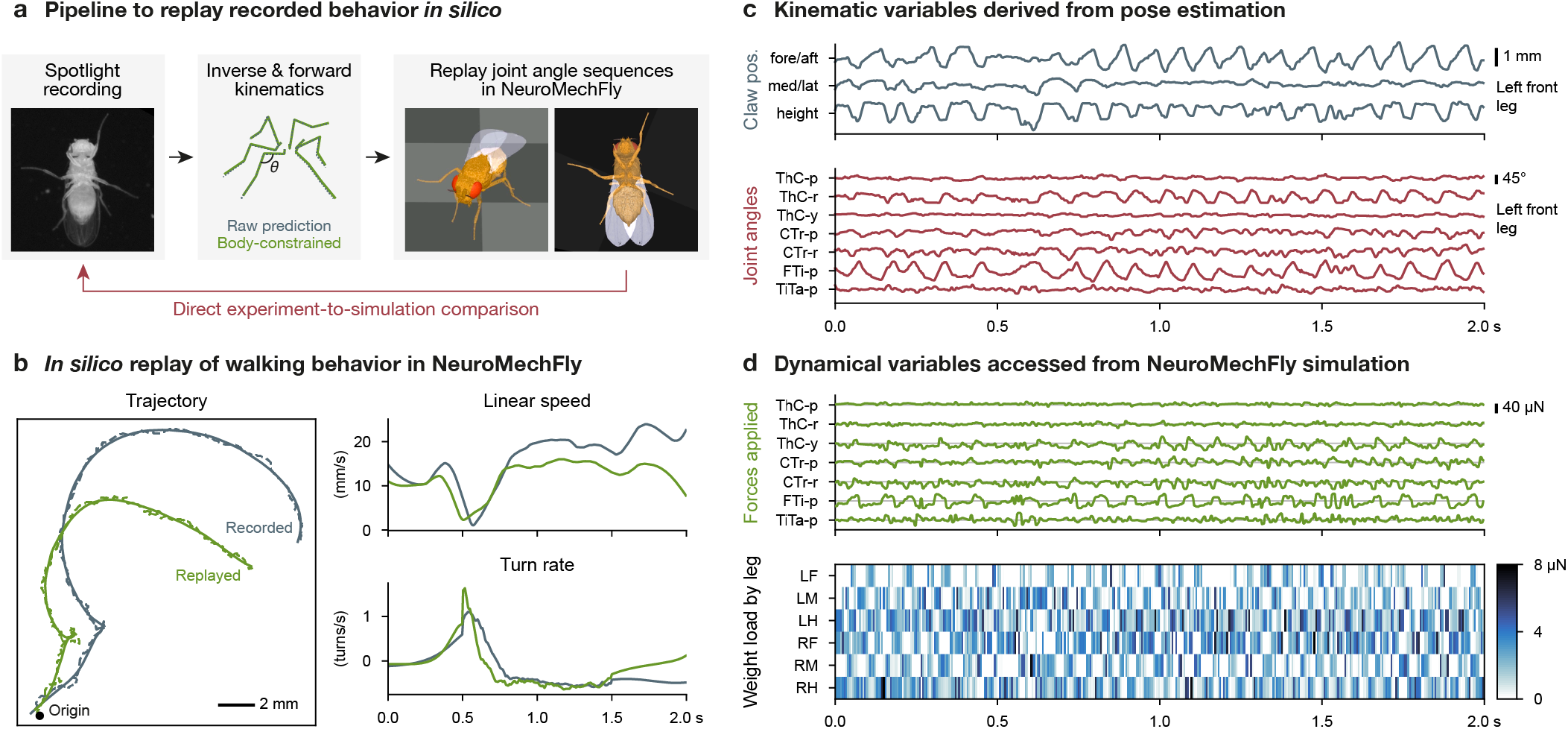
Kinematics derived from *Spotlight* recordings and *PoseForge* are replayed in a biomechanical simulation to infer unmeasured dynamical variables. **(a)** Keypoints are extracted from *Spotlight* images using *PoseForge*. Joint angles are reconstructed via inverse kinematics and used to actuate the biomechanical model, NeuroMechFly. Replay of experimentally inferred joint angle trajectories produces simulated kinematics directly comparable with experimental measurements. Some level of mismatch is expected, as generating forces to track recorded kinematics inevitably introduces a delay. **(b)** Example walking sequence consisting of straight walking followed by a rapid turn and subsequent curved walking. (left) Trajectory of the real fly (blue) and the NeuroMechFly model (orange) during replay of experimentally recorded joint angles. Dashed and solid lines indicate raw and temporally smoothed trajectories, respectively. (top right) Linear speed and (bottom right) turning rate of the real and simulated fly during the same time window (after smoothing). **(c)** Left front leg kinematic variables derived from *PoseForge* and subsequent inverse/forward kinematics. (top) Egocentric 3D claw position. (bottom) Joint angles (ThC: thorax-coxa; CTr: coxa–femur; FTi: femur–tibia; TiTa: tibia–tarsus; p: pitch; r: roll; y: yaw). Time window matches that in panel b. **(d)** Dynamical variables inferred from simulation. (top) Left front leg joint torques required to track real kinematics. (bottom) Load distribution across all legs during locomotion. The simulated fly weighs 10 µN ^66^.

We then simulated an untethered NeuroMechFly model walking on a flat ground plane, driven by position actuators that track reconstructed joint angles (**Figure 3c**). Unlike prior literature that used deep reinforcement learning to train controllers to match recorded behavior ^19^, our approach relies on joint angles alone and requires no training. The simulated fly reproduced speed and turning rates that closely matched real-animal kinematics, including sharp in-place turns (saccades), although global trajectories diverge over time due to accumulated error (**Figure 3b**). This indirectly validates the quality of pose estimation and inverse kinematics, and indicates that NeuroMechFly captures key aspects of the physics relevant to locomotion. During replay, we recorded joint torques and ground reaction forces across all legs (**Figure 3d**), providing a complementary dynamical interpretation of behaviors recorded in *Spotlight*. Because inferred dynamics depend on biomechanical parameters that are difficult to measure experimentally, we performed sensitivity analyses by varying physics parameters. Although absolute magnitudes differ, patterns were largely preserved (**Extended Data Fig. 4**), suggesting that *in silico* replay can estimate latent dynamical variables even when exact physical parameters are uncertain.

### Measuring muscle activity in freely behaving flies from Spotlight recordings

Although replaying kinematic recordings *in silico* enables scalable *inference* of biomechanical variables, these simulations do not directly measure the underlying physiological activity. This motivated us to develop direct muscle imaging capabilities in *Spotlight*. To demonstrate this capability, we recorded the activities of long-tendon muscles in the legs. These are physically located in the femur but control distal claw movements through a tendon ^67^.

We observed that these muscles tended to become active upon manual perturbation of the arena, possibly to maintain a stable posture. To test this hypothesis, we delivered brief pulses of vibration using a small motor attached to the behavior arena. Actuating the motor at regular intervals elicited a reliable increase in long-tendon muscle activity that remained sustained throughout the stimulation period (**Figure 4a**). Notably, mechanical stimulation produced little to no visible movement of the animal (**Video 7**), indicating that the observed fluorescence changes were not due to motion artifacts. In line with this, fluorescence traces extracted from GFP control animals—whose fluorescence levels are independent of muscle activity—remained largely constant during vibration (**Figure 4b**).

**Figure 4:**
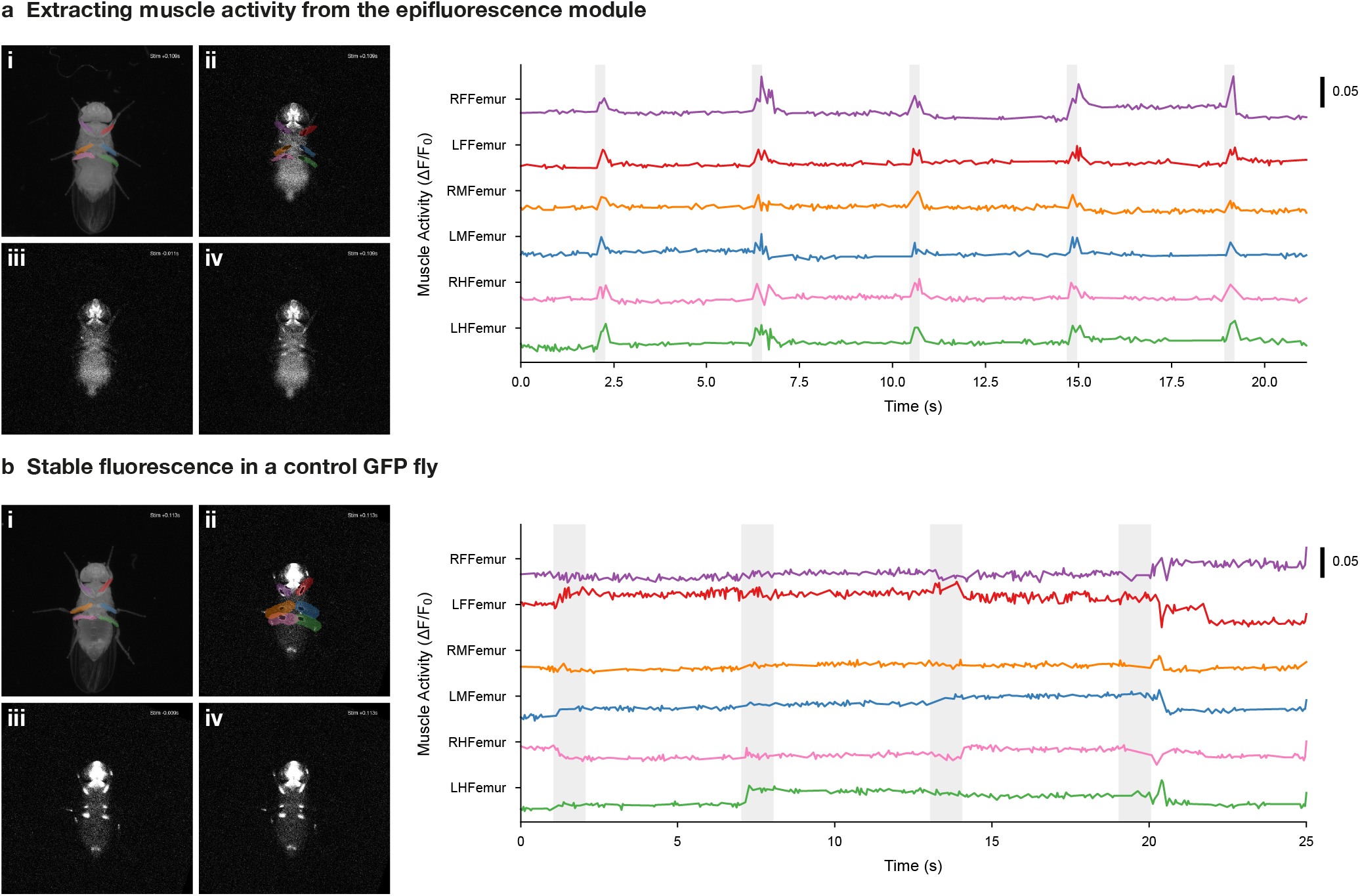
Optically recording and extracting the activity of long-tendon muscles using Spotlight and Poseforge. Fluorescence traces acquired by measuring either **(a)** the calcium indicator GCaMP8m-RSET, or **(b)** GFP during mechanical vibration stimulation periods (light grey shading). Behavioral and muscle imaging data were acquired synchronously. **(i)** Segment regions of interest (ROIs) were generated from behavioral images using *PoseForge* and **(ii)** mapped to the muscle imaging data using a custom homography transformation. ROIs were denoised and dilated, and the 500 brightest pixels within each ROI were selected (black). When the muscle was active, these pixels cluster along the muscle, whereas during inactivity they are more spatially dispersed. **(iii)** Normalized muscle fluorescence images immediately before or **(iv)** a few milliseconds after mechanical stimulation. See also **Extended Data Fig. 5** for fluorescence traces from more flies and with multiple vibration pulses overlaid.

Although conclusive characterization of such behavioral control strategies will require additional experiments both with our system and using other approaches, this simple proof of concept illustrates how whole-body, large-scale measurements in minimally restricted settings can expose patterns that would likely be missed in more targeted experiments, thereby providing an entry point for hypothesis generation. Moreover, because anatomical links between muscles, motor neurons, and central nervous system neurons have been systematically characterized in *Drosophila* ^68,69^, muscle activity measurements can potentially provide a bridge between behavioral observations and the underlying motor circuits through connectomic resources ^15,16,17^.

## Discussion

Here we have presented an end-to-end framework that integrates closed-loop experimentation, synthetic-data–driven pose estimation, and biomechanical simulation to study natural behavior in freely behaving flies. The resulting multi-level dataset includes observable kinematics, muscle activity across limbs, and inferred dynamical variables, enabling quantitative analyses across kinematic and mechanical domains, both *in vivo* and *in silico*. High-resolution measurements of unconstrained behavior are broadly valuable in systems neuroscience; in *Drosophila* in particular, advances in connectomics ^14,15,16,17^ and whole-body modeling ^18,19,20^ drive the demand for detailed behavioral ground truth data to constrain and validate connectome-constrained simulations.

Althougb the present work focuses on muscle activity as a readout of motor output, the approach developed here could also be extended to neural recording. Untethered brain imaging in freely moving flies (with dissection) has been demonstrated using the Flyception system ^7,8^. In the future, incorporating a similar neural imaging configuration in *Spotlight* could combine high-resolution behavioral quantification with neural activity measurements, providing a powerful framework for linking neural circuit dynamics to motor output and behavior.

Although developed for *Drosophila*, the general strategies in our system extend beyond flies. First, closed-loop tracking videography mitigates the general trade-off between spatiotemporal resolution and unconstrained behavior. Second, our annotation-free kinematic analysis pipeline shows how anatomically accurate body models, paired with advances in generative AI, can scale behavioral quantification, both in data volume and in richness of information extracted. Similarly, self-supervised learning further reduces the reliance on manual human annotations by bootstrapping behavior representations directly from data that can be scalably generated. Finally, integration with NeuroMechFly illustrates how biomechanical digital twins—and, in future extensions, *neuro*mechanical digital twins—can infer latent quantities and bridge experimental measurement with computational modeling.

Several limitations and future directions remain. First, continuous blue light excitation for muscle imaging can agitate animals, leading to fast but atypical walking. This may be mitigated by interleaving recording and recovery periods. Second, synthetic image generation was trained using kinematics derived from tethered recordings, imposing a biased prior that reduces generalization to behaviors unrepresented in the reference dataset. This can be mitigated through iterative bootstrapping: namely using untethered kinematics extracted from *Spotlight* and *PoseForge* to expand the generator’s training pool, and retraining the pose models that follow. Third, current models operate on full-body images, which, at fixed model size, limits the effective spatial resolution and prevents finer muscle segmentation. Larger models or patch-based pipelines can increase the effective resolution in future implementations. Fourth, fidelity of both synthetic image generation and *in silico* replay depends on the rigging and parameterization of the NeuroMechFly model (e.g., precise joint attachment points and rotation axes, passive joint properties, and contact dynamics), which would benefit from further experimental calibration. Finally, although muscle calcium imaging is non-invasive and scalable, extraction and quantitative interpretation of fluorescence traces will require continued development of downstream analysis and validation pipelines.

Despite these limitations, this work demonstrates how advances in engineering, generative AI, and biomechanical simulation can be combined into an integrated experimental–computational workflow. Such co-designed systems enable quantitative studies under technically demanding conditions that more closely reflect natural behavior, supporting mechanistic links between neural circuits, biomechanics, and behavior.

## Methods

### *Spotlight* design and construction

The main components of *Spotlight* include a behavior recording system, a muscle imaging system, motorized motion stages, electronic controllers, and a customizable arena in which flies can move freely. *Behavior recording*. Fly behavior was recorded using a JAI SP-5000M-CXP4 5-megapixel camera (JAI A/S, Denmark) with a Laowa 100mm f/2.8 2x Ultra Macro APO lens (Venus Optics, China). Image data were acquired using a Euresys Coaxlink Quad G3 frame grabber (EURESYS S.A., Belgium) via a CXP-6 CoaXPress cable (STEMMER IMAGING AG, Germany). Infrared illumination was provided using a 850 nm CCS LDR2-74IR2-850-LA ring light controlled by a CCS PD3-3024-3-EI light controller (OPTEX GROUP Co., Ltd., Japan). In a typical experimental configuration, the magnification level of the lens was set to ~ 0.9:1 and the aperture was set to ~ f/8.

#### Muscle recording

Muscle fluorescence was recorded using a pco.panda 4.2 sCMOS USB camera (Excelitas Technologies Corp., USA) with a second Laowa 100mm f/2.8 2x Ultra Macro APO lens (Venus Optics, China). Fluorescence excitation was provided using a 470 nm Thorlabs M470L3 LED light driven by a Thorlabs LEDD1B driver (Thorlabs Inc., USA). The excitation beam profile was shaped using Thorlabs AC254-050-AB and LA1509-ML lenses. In a typical experimental configuration, the magnification level of the lens was set to ~ 1.2:1 and the aperture was set to ~ f/5.6.

#### Optical path separation

The optical filter configuration is shown in **Extended Data Fig. 1a**. To record behavior, a Semrock FF01-715/LP-25 long-pass filter placed between the behavior camera and its lens isolated the infrared behavior recording path from the muscle imaging path. For muscle fluorescence imaging, a Semrock FF01-466/40-25 band-pass filter positioned in front of the blue LED served as the excitation filter, and a Semrock FF03-525/50-25 band-pass filter placed between the muscle camera and its lens served as an emission filter. A Semrock Di02-R488-25x36 long-pass dichroic mirror reflected blue excitation light toward the fly while transmitting the green fluorescence emission and infrared light toward the cameras. A second Semrock FF775-Di01-25x36 long-pass dichroic mirror reflected green emission light toward the muscle camera while transmitting infrared light toward the behavior camera. The two dichroic mirrors were mounted with a 90^°^ rotation about the vertical axis to allow the muscle camera and the blue excitation LED to be installed on different sides of the behavior camera (**Extended Data Fig. 1a**). All Semrock filters were supplied by IDEX Health & Science LLC (USA).

#### Behavior arena

The behavioral arena consisted of two 1 mm-thick glass slides separated by 2 mm-thick walls, creating a rectangular open-space of 72 × 48 mm (**Extended Data Fig. 1b**). Alternative arena configurations can easily be made by laser cutting different wall patterns in 2 mm or by using thicker acrylic plates. The arena was sandwiched between two 2 mm acrylic frames and clasped with two thin clamps, minimizing the clearance distance between the moving illumination LED ring and the glass floor (~ 3 mm thickness). Flies walked on both the floor and ceiling of these arenas, but full-leg images on *Spotlight* could only be obtained when the fly was on the floor. To facilitate fast flipping of the chamber when flies walked on the ceiling (and rapid loading of flies), the arena-holder was up-down symmetric, and was held in place by simply dropping it into an opening in a suspended acrylic slide with notches matching those in the arena-holder frames, ending up flush with the base of the arena-holder. Microscope slides were used for the floor and ceiling of the arena for optical clarity (Ted Pella 260240-4 ClariTex Super Mega Slides, Plain; Ted Pella, Inc., USA), while other parts of the arena were laser-cut from generic acrylic plates (PMMA Acrylglas; Amsler & Frey AG, Switzerland).

#### Optional optogenetic stimulation or silencing

A flat LED panel (CCS TH2-51x51RD, OPTEX GROUP Co., Ltd., Japan) can be installed to provide red-light optogenetic stimulation via activation of CsChrimson ^47^ (**Extended Data Fig. 1c**). Alternatively, a blue LED (e.g., CCS TH2-51x51BL) can be used for optogenetic silencing via activation of GtACR ^48^. Although this configuration is incompatible with GCaMP-based muscle imaging. The LED was mounted on a support arm attached to the muscle camera, allowing the stimulation light to track the animal. An aperture of adjustable size was placed in front of the LED to localize illumination and reduce heating. The LED was driven by the same CCS light controller. A short-pass filter (Edmund Optics 54-516; Edmund Optics BV, The Netherlands) blocked infrared wavelengths to avoid affecting behavioral recordings.

#### Motion stages

Two stacked Zaber LSQ150A-E01CT3A motorized linear stages, controlled by a Zaber X-MCC2 controller (Zaber Technologies Inc., USA), provided actuation along the *x* and *y* axes. The behavior and muscle camera assemblies were mounted on this actuated stage.

#### Mechanical construction

The structure holding the two dichroic mirrors, a rack used to align the behavior camera, and the clamps and alignment pins of the behavior arena were custom-designed and 3D-printed. An extension plate for mounting the muscle camera assembly to the actuated stage was custom-designed and CNC-manufactured. The plate holding the infrared ring light, the arena mounting plates, and the arena itself were laser-cut from acrylic plates. Counterbore holes were drilled into a Thorlabs MB2020/M breadboard to allow mounting onto the motion stages. The muscle camera was mounted on a Siskiyou AB-U (metric) bracket (Siskiyou Corporation, USA). All remaining components were standard optomechanical parts from Thorlabs Inc., USA. CAD files for all custom components, a complete assembly model, and a full parts list are available at https://go.epfl.ch/spotlight-poseforge.

### Spatial and cross-camera calibration

Operating Spotlight requires calibration between four coordinate systems: the *physical* coordinates (position in the arena in mm, ***ω***_*p*_ = [*x*_phy_, *y*_phy_]), the *stage* coordinates (positions of the motion stages in mm, ***ω***_*s*_ = [*ω*_stage_, *y*_stage_]), the behavior camera *pixel* coordinates (location on the acquired image in pixels, ***φ***_*b*_ = [*i*_beh_, *j*_beh_]), and the muscle camera *pixel* coordinates (***φ***_*m*_ = [*i*_mus_, *j*_mus_])).

The cameras were first approximately aligned using a printed target. To estimate the transformation between the coordinate systems, we placed a printed board with ArUco markers ^70^ on top of the arena. The arena was then scanned using the motion stages, pausing at 2 mm intervals and acquiring an image from both cameras at each stop. Corners of the ArUco markers were detected in pixel coordinates and paired with the corresponding physical coordinates defined by the known ArUco marker layout.

Assuming affine transformations between coordinate systems, we fitted vectors ***β***_∗_ ∈ ℝ^5×1^ using the following linear models:

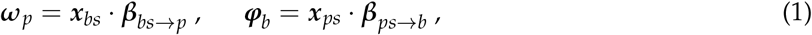

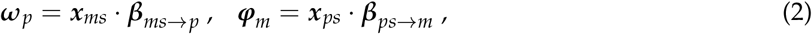

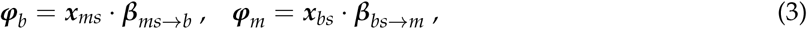

where the input vectors are defined as

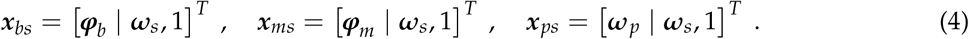

These transformations were used both for real-time tracking during experiments and for postprocessing after the experiment.

### Closed-loop control and recording

Images acquired by the cameras were transmitted to a recording computer (Intel® Core™ i7-11700K processor; 4 × 16 GB Kingston 9905743-058.A00G DDR4 RAM; CORSAIR Force Series™ MP510 4 TB M.2 SSD for data storage and Samsung 500 GB 970 EVO Plus M.2 SSD for the operating system; no dedicated graphics card; operating on Ubuntu 24.04.4 LTS). Behavior images were acquired using the Euresys eGrabber C++ acquisition library; muscle images were acquired using the pco.cpp C++ Software Development Kit (SDK). Raw images were compressed and written to disk using eight parallel threads (four for behavior images and four for muscle images). Muscle images were stored as 16-bit TIFF files to preserve dynamic range. Behavior images were stored as three-channel 8-bit JPEG files for efficient compression: each file contained three sequential frames encoded as separate channels, leveraging the high compression ratio of the JPEG format for similar single-channel images. Motion stage positions were read at 75 Hz and saved using an additional thread.

In parallel, a separate thread estimated the centroid position of the fly in behavior images at 30 Hz, independently of the recording frame rate. The fly was detected by binarizing the image using a threshold of 75 (out of 255) and selecting the largest connected component with a minimum size of 1,000 pixels. When arena boundaries were visible, pixels within 0.5 mm of the arena boundaries were cropped to remove bright reflections from the edges of the acrylic plates. The centroid of the detected component was computed and converted from pixel coordinates to physical coordinates using the fitted transformations based on the current motion stage positions. If the resulting physical coordinates differed from current motion stage positions by more than 0.1 mm, the stages were set to move to the fly centroid position using position control at speeds of up to 63 mm s^−1^.

All cameras and illumination sources were hardware-triggered to ensure synchronization. The recording computer sent control signals to an Arduino Nano ESP32 microcontroller (240 MHz clock speed; Arduino S.r.l., Italy, and Espressif Systems, China), which in turn output TTL signals to the Euresys frame grabber controlling the behavior camera, the PCO camera, the CCS light controller for behavior illumination, and the Thorlabs light controller for muscle excitation. When optogenetic stimulation was used, the microcontroller also controlled the stimulation light source.

All cameras and lights were hardware-triggered to ensure synchronization. The recording computer sent control signals to an Arduino Nano ESP32 microcontroller (240 MHz clock speed; Arduino S.r.l., Italy, and Espressif Systems, China), which in turn generated TTL signals to the Euresys frame grabber controlling the behavior camera, the PCO camera directly, the CCS light controller for behavior illumination, and the Thorlabs light controller for muscle excitation. When optogenetic stimulation was used, the microcontroller also controlled the stimulation LED.

A graphical user interface ran in a separate thread and provided a live preview of the camera streams and motion stage positions. The interface allowed the user to start and stop recordings and adjust acquisition parameters.

After the experiment, a postprocessing pipeline converted behavior images from JPEG files into a more efficiently compressed video format (H.264 codec). Motion stage positions were interpolated to match the timestamps of behavior frames. Muscle images were warped using the previously fitted affine transforms to match behavior images. A simple 2D pose estimation model was trained using SLEAP ^25^ to track the neck, thorax center, and abdomen tip of the animal. From these keypoints, the body center and heading of the fly were estimated, and frames were rotated and cropped to a smaller 900 × 900 pixel bounding box centered on the fly, with the animal oriented upward. This postprocessing pipeline was parallelized across multiple processes.

### Translating simulation renderings to the appearance of *Spotlight* recordings

Each synthetic frame was independently generated. Although this approach reduced temporal continuity, it minimized the risk that downstream pose estimation models exploited temporally correlated artifacts arising from the biased prior of tethered kinematic data when applied to freely behaving animals. We used Contrastive Unpaired Translation (CUT) ^29^ to translate NeuroMechFly renderings into *Spotlight*-like images. The CUT architecture consists of a generator that performs image translation, a discriminator that acts as a binary classifier distinguishing synthetic outputs from real images in the training dataset, and a contrastive projection head.

Briefly, image patches—in both simulation and *Spotlight* domains—are mapped to latent representations by the projection head. An InfoNCE loss ^59^ encourages latent vectors corresponding to the same spatial location in the input (simulation domain) and translated (*Spotlight* domain) images to be similar, while enforcing dissimilarity between vectors extracted from different spatial locations or from different images, thereby promoting preservation of spatial pose features. In parallel, an adversarial loss ^58^ encourages the discriminator to correctly classify real and synthetic samples, thereby training the generator to produce images that are indistinguishable from real experimental recordings. By combining the patch-based InfoNCE loss with the adversarial loss, the model learns to generate realistic images while preserving pose information from the NeuroMechFly simulations.

The generator was implemented using a StyleGAN ^71^ architecture, while the discriminator used a PatchGAN ^72^ architecture. We trained 31 variants of the model with different hyperparameter settings (**Table 1**). From these, we manually selected the eight best models at epochs that produced the best qualitative results, as the two partially competing terms of the overall loss made quantitative selection of the best model difficult. Key remaining hyperparameters that were not tuned included the PatchGAN receptive field in the discriminator (70 × 70 pixels), the number of filters in the final convolutional layer of the discriminator (64), maximum number of training epochs (200), the optimizer (Adam ^73^), the learning rate (0.0002, decaying linearly to 0 over the last 100 epochs), the Adam momentum weights (*β*_1_ = 0.5, *β*_2_ = 0.999), and the adversarial loss (least-square GAN ^74^). These choices largely follow the default values in the CUT implementation ^29^. The training dataset consisted of 10,000 simulated images and 10,000 experimentally recorded images sampled at random. The testing dataset consisted of 1,000 simulated images and 1,000 recorded images, randomly sampled from trials not included in the training dataset. Training was performed on the EPFL SCITAS Kuma cluster; each training job was allocated one NVIDIA® L40S GPU (48 GB), 64 GB of RAM, and 32 threads on an AMD EPYC™ 9334 CPU.

**Table 1:** Hyperparameters for synthetic data generation model. “Gen. #blocks”: number of StyleGAN blocks in the generator; “Gen. #filters”: number of filters in the last convolution layer in the generator; “GAN loss weight”: weight of the adversarial loss relative to the InfoNCE loss; “Batch size”: number of samples per training batch; “Selected epoch”: training epoch that produces the best qualitative results (only if the model is selected for pose estimation training).

| Trial | Gen. #blocks | Gen. #filters | GAN loss weight | Batch size | Selected epoch |
| --- | --- | --- | --- | --- | --- |
| 1 | 2 | 16 | 0.2 | 2 | 121 |
| 2 | 2 | 16 | 0.5 | 2 |  |
| 3 | 2 | 16 | 1.0 | 2 |  |
| 4 | 2 | 32 | 0.2 | 2 |  |
| 5 | 2 | 32 | 0.5 | 2 |  |
| 6 | 2 | 32 | 0.5 | 8 |  |
| 7 | 2 | 32 | 1.0 | 2 |  |
| 8 | 2 | 48 | 0.2 | 2 |  |
| 9 | 2 | 48 | 0.5 | 2 |  |
| 10 | 2 | 48 | 1.0 | 2 |  |
| 11 | 2 | 64 | 0.5 | 8 |  |
| 12 | 6 | 16 | 0.1 | 2 |  |
| 13 | 6 | 16 | 0.1 | 4 |  |
| 14 | 6 | 16 | 0.2 | 2 |  |
| 15 | 6 | 16 | 0.2 | 4 | 200 |
| 16 | 6 | 32 | 0.1 | 2 |  |
| 17 | 6 | 32 | 0.1 | 4 | 161 |
| 18 | 6 | 32 | 0.2 | 2 |  |
| 19 | 6 | 32 | 0.2 | 4 |  |
| 20 | 6 | 32 | 0.5 | 2 | 161 |
| 21 | 6 | 32 | 0.5 | 4 | 141 |
| 22 | 6 | 32 | 1.0 | 2 |  |
| 23 | 6 | 32 | 1.0 | 4 | 161 |
| 24 | 6 | 48 | 0.1 | 2 | 141 |
| 25 | 6 | 48 | 0.1 | 4 | 141 |
| 26 | 6 | 48 | 0.2 | 2 |  |
| 27 | 6 | 48 | 0.2 | 4 |  |
| 28 | 6 | 64 | 0.5 | 2 |  |
| 29 | 6 | 64 | 0.5 | 4 |  |
| 30 | 6 | 64 | 1.0 | 2 |  |
| 31 | 6 | 64 | 1.0 | 4 |  |

### Contrastive pretraining

Let ***x***_*ϕ,i*_ denote the *ϕ*-th synthetic variant of the *i*-th frame. Let Enc be a shared feature extractor and Proj a contrastive projection head (both implemented as neural networks). The latent representation of ***x***_*ϕ,i*_ in the feature space is given by ***h***_*ϕ,i*_ = Enc(***x***_*ϕ,i*_), and the representation in the projection space is ***z***_*ϕ,i*_ = Proj(***h***_*ϕ,i*_). Contrastive similarity is defined in the ***z***-space, while the ***h***-space representations are reused for downstream pose estimation tasks. Following InfoNCE ^59^ and SimCLR ^61^, the similarity function *f* between two samples is defined as

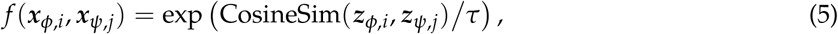

where a positive “temperature” constant *τ* controls the sensitivity of *f* to similarities and dissimilarities. Empirically, we set *τ* = 0.1. We then used an MIL-NCE loss ^62^ for contrastive learning:

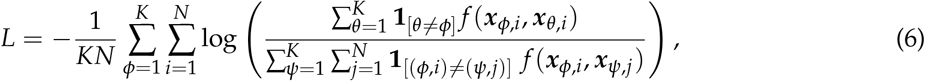

where **1**_[*c*]_ equals 1 when the condition *c* holds and 0 otherwise.

In practice, the loss was estimated through Monte Carlo sampling via gradient descent. Efficient implementation required random access to frames from different videos within each batch. However, this was inefficient whether frames were encoded as videos (random access requires decoding preceding frames) or as independent images (which leads to large storage overhead). To address this issue, we partitioned the dataset into small “atomic batches” and stored them sequentially as videos. At each training iteration, a random set of atomic batches was sampled, but the contents of each atomic batch remained fixed. Because the frames within an atomic batch were always loaded together, they could be read sequentially and data loading could be parallelized across multiple atomic batches. Empirically, the dataset was partitioned into ~ 116,000 atomic batches, each containing *n* = 32 frames with *k* = 4 variants per frame. To ensure that sampled frames were temporally distinct, frames were selected with a gap of 60 frames (0.2 s) between successive samples. During training, *b* = 30 atomic batches were sampled per iteration.

The loss was implemented by first flattening the input tensor from (*b, n, k*, …) to (*bnk*, …), where “…” denotes the image dimensions. Passing the batch through Enc and Proj yielded feature vectors ***Z*** ∈ ℝ^*bnk*×*w*^, where *w* is the dimensionality of the projection space. Pairwise cosine similarities between all samples are then computed, forming a similarity matrix ***S*** ∈ ℝ^*bnk*×*bnk*^. For each combination of indices *β* ∈ [1, *b*], *ν* ∈ [1, *n*], *κ* ∈ [1, *k*], the corresponding flattened row index is *u* = *βnk* + *νk* + *κ*. Then, for each row ***s***_*u*_ = [*s*_*u*,1_, …, *s*_*u,bnk*_], the elements are grouped into three categories. First, the diagonal element *s*_*u,u*_ represents the similarity of the sample with itself and is discarded. Second, elements with column indices {*βnk* + *νk* + *κ*′ | *κ*′ ∈ [1, *k*], *κ*′/= *κ*} correspond to similarities between different variants of the same frame. We denote these by vector ***s***^+^ ∈ ℝ^*k*−1^. Third, all remaining elements correspond to similarities between different frames, which we denote by vector 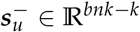. The loss for each row ***s***_*u*_ is then

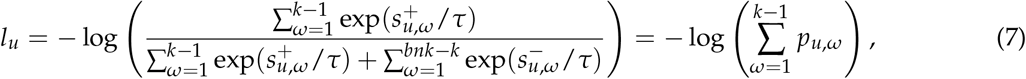

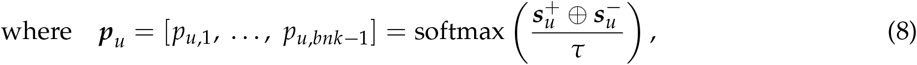

where ⊕ denotes vector concatenation. For each iteration, the loss in **Equation 6** is estimated by averaging *l*_*u*_ over *u* = 1 … (*bnk*). In practice, this loss was minimized using the optimizer ^73^ with a learning rate of 3 × 10^−4^ and *L*_2_ weight decay of 10^−4^.

The feature extractor Enc was implemented as a pretrained ResNet-18^63^ (TorchVision “ResNet18_ Weights.IMAGENET1K_V1” ^75^). Each input image ***x*** was resized to 256 × 256 pixels. Because the pretrained network expects RGB inputs, the single grayscale channel was repeated across three channels. The final pooling, fully connected, and softmax layers of ResNet-18 were removed, resulting in a 512-channel, 8 × 8 feature map (the latent representation ***h***). The contrastive projection head Proj was implemented as a three-layer multilayer perceptron (input size 512, hidden size 512, output size 256). Hidden layers used ReLU activation, and the output layer had no nonlinear activation. Before entering the projection head, ***h*** was average-pooled to size (512, 1, 1) and flattened. Similarities were computed in the resulting 256-dimensional projection space where ***z*** resides.

### Predicting 3D keypoint positions

The output feature map of Enc is successively upsampled using a decoder Up with skip connections from the corresponding layers in Enc (following the U-Net architecture ^64^). We used four upsampling blocks, each consisting of a transposed convolution layer, a concatenation convolution layer, batch normalization, another convolution layer, and ReLU activation. In each upsampling block, the transposed convolution layer receives a *c*_in_-channel, *s*_in_ × *s*_in_ feature map and outputs a *c*_in_-channel, 2*s*_in_ × 2*s*_in_ feature map using a kernel size of 2 and a stride of 2. The concatenation convolution layer receives the output of the transposed convolution layer and a *c*_copy_-channel, 2*s*_in_ × 2*s*_in_ feature map copied from the corresponding downsampling block in Enc (the U-Net skip connection). The concatenation convolution layer outputs a *c*_out_-channel, 2*s*_in_ × 2*s*_in_ feature map using a kernel size of 3 and padding of 1. This output is batch-normalized, transformed using a ReLU activation function, and then passed through a final convolution layer that preserves the spatial dimensions using a kernel size of 3 and a padding of 1, followed by batch normalization and ReLU activation. The four upsampling blocks are configured with *c*_in_ = 512, 256, 128, 64, *c*_copy_ = 256, 128, 64, 64, *c*_out_ = 256, 128, 64, 64, *s*_in_ = 8, 16, 32, 64. This produces a 64-channel, 128 × 128 feature map ***u***.

The upsampled feature map ***u*** is then given as input to two separate decoding heads: H_*xy*_, which predicts 2D heatmaps for keypoint positions in row-column coordinates, and H_z_, which predicts distributions over keypoint depths (distance from the camera). The decoding head H_*xy*_ consists of a single convolution layer with kernel size 3 that transforms the 64-channel input ***u*** into a 32-channel output ***v***_2d_ (one channel per tracked keypoint: thorax-coxa, coxa-trochanter, femur-tibia, tibia-tarsus, and claw for each leg, plus two antennae). The spatial resolution remains 128×128 pixels. The probability of the keypoint falling into each heatmap bin ***p***_2d_ is then obtained from logits ***v***_2d_ using softmax with temperature *τ* = 0.8: ***p***_2d_ = softmax(***v***_2d_/*τ*). The final predictions for the row and column coordinates for each keypoint *a* (denoted by 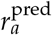,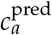) are computed using a “soft argmax” function:

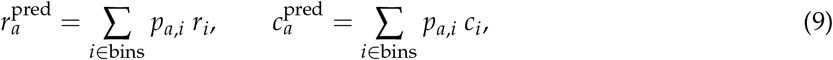

where *p*_*a,i*_ is the probability assigned to bin *i*, and *r*_*i*_, *c*_*i*_ are the row and column coordinates of the center of bin *i*.

Furthermore, we quantified prediction confidence *γ*_2d_ ∈ ℝ^32^ (one value per keypoint) based on the entropy of the predicted distribution:

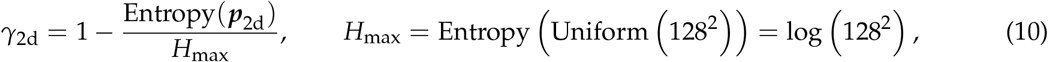

where Entropy(·) is the information entropy of each categorical distribution. The maximum entropy possible *H*_max_ is that of a uniform distribution, which evaluates to log(*n*) for a distribution over *n* = 128^2^ bins.

To predict keypoint depth, the range [−2, 2] mm relative to the arena floor is first discretized into 64 evenly spaced bins. This allows us to predict probabilities of the keypoint falling into each of these bins, similar to a 2D heatmap. In the H_z_, ***u*** is first average-pooled to a 1 × 1 feature map and flattened. A convolution layer maps the 64-channel input to 128 channels, followed by group normalization with 32 groups. The resulting feature map is flattened and passed through a fully connected layer that maps the 128-dimensional input to a 2048-dimensional output. This output is reshaped into logits ***v***_depth_ ∈ ℝ^32×64^, where 32 is the number of keypoints and 64 is the number of depth bins.

Similar to H_*xy*_, our temperature-regulated probabilities are given by ***p***_depth_ = Softmax(***v***_depth_/*τ*), where *τ* = 0.8. The final depth prediction 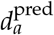 for each keypoint *a* is

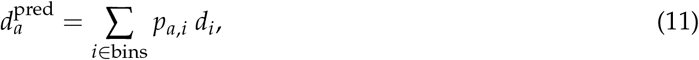

where *d*_*i*_ is the center of depth bin *i*. Prediction confidence is quantified by

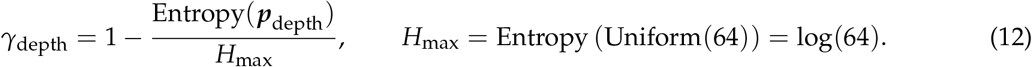

Predicting row–column coordinates and depth in separate pathways improves memory and parameter efficiency. Instead of predicting probabilities in a full 3D voxel grid, the model predicts a 2D heatmap and a 1D depth distribution. Although this factorization removes explicit coupling between spatial and depth coordinates, the two predictions remain implicitly coupled through the shared feature representation ***u*** produced by Up (the shared pathway until ***u*** accounts for ~ 98% of model weights, **??**).

Prior to training, the ground-truth *x*-*y*-*z* positions obtained from NeuroMechFly simulations were transformed into row-column-depth coordinates. To train the Up ⊕ H_*xy*_ pathway (where ⊕ denotes concatenation), we generated ground-truth distributions ***q***_2d_ centered on the true row-column positions using a 2D Gaussian with a standard deviation of 2 pixels on a 128×128 grid (matching the size of the output of H_*xy*_, but with a stride of 2 compared to the 256×256-pixel input). The loss is then

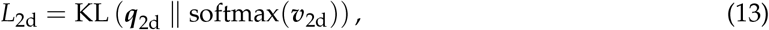

where KL(· ∥ ·) denotes the Kullback–Leibler divergence.

The Up ⊕ H_z_ pathway was trained using two loss terms. The first was a cross-entropy (CE) loss treating depth prediction as classification over discrete bins:

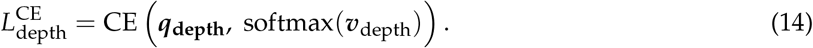

The second term was an *L*_1_ regression loss:

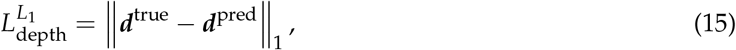

where ***d***^pred^, ***d***^true^ are the predicted and true depth vectors.

We empirically defined the final loss as 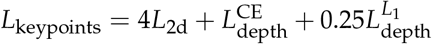 decoders Up, H_*xy*_, and H_z_ using this loss. Throughout the process, the 2D and depth pathways shared the same instantiation of the encoder Enc. We trained the models using an Adam optimizer ^73^. To primarily update the upsampling core Up and the decoding heads H_*xy*_, H_*z*_ while more conservatively preserving the pretrained feature extractor Enc, we used a learning rate of 3 × 10^−4^ for Up, H_*xy*_, H_*z*_, and a much lower learning rate of 3 × 10^−5^ for Enc. The weight decay parameter was set to 10^−5^.

### Predicting body segmentation maps

To predict body segmentation maps, we defined 28 anatomical segments: the coxa, trochanter-femur, tibia, and tarsus for each leg (6 × 4=24), left and right antenna (2), thorax (1), and all remaining body parts (1). Together with the background class, these constitute *m* = 29 classes that each pixel may belong to. The segmentation task is therefore formulated as a per-pixel *m*-class classification problem.

As in the 3D keypoint model, the pretrained feature extractor Enc was used to compute a feature map ***h*** from the input image ***x***. A segmentation decoding head H_seg_ then upsamples ***h*** to a 29-channel, 256 × 256 output ***v***_seg_, whose elements represent the logits for each pixel belonging to each class. Internally, the first part of H_seg_ shared the same architecture as Up—successively upsampling the feature map while incorporating U-Net-style skip connections from the downsampling pathway of Enc. Once we reach the 64-channel, 128 × 128 feature map ***u***, it is upsampled once more using a transposed convolution layer with kernel size 2 and stride 2, generating a 32-channel, 256 × 256-pixel feature map matching the input resolution. Finally, a linear layer transforms the 32-dimensional final feature vector to a 29-dimensional logits vector on a per-pixel basis (this is actually implemented as a convolution layer with kernel size 1, stride 1, input channels 32, and output channels 29). For each pixel *i*, we denote the vector of logits [*v*_1,*i*_, …, *v*_*m,i*_] by ***v***_seg,*i*_ ∈ ℝ^*m*^. We can then convert these logits to probabilities ***p***_seg,*i*_ = [*p*_1,*i*_, …, *p*_*m,i*_] = softmax(***v***_seg,*i*_).

Similar to the 2D keypoint heatmap, the confidence *γ*_seg_ of our prediction can be quantified using the normalized entropy of the predicted distribution:

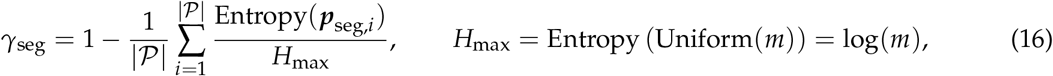

where 𝒫 is the set of all pixels in the image.

The Enc ⊕ H_seg_ pathway was trained using a combination of Dice loss and cross-entropy loss. Intuitively, the Dice loss treats segmentation as *m* independent mask prediction tasks, emphasizing the *spatial* consistency of each class mask while ignoring distribution across classes. In contrast, the cross-entropy loss treats segmentation as 256 × 256 independent *classification* tasks, emphasizing *per-pixel* accuracy across distinct classes but handling different pixels independently.

Let *e*_*c,i*_ denote the ground-truth label for class *c* at pixel *i* (1 if the pixel belongs to the corresponding class and 0 otherwise). For each class *c*, the Dice loss is

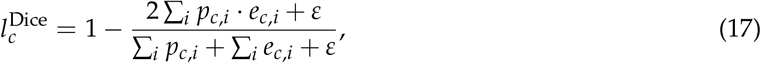

where *ε* = 10^−6^ is added for numerical stability. The total Dice loss 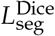 is obtained by averaging 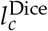 over classes and training samples.

The cross-entropy loss for each pixel *i* is

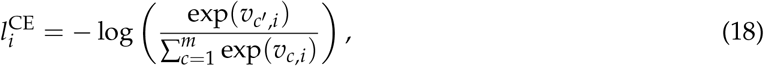

where *c*′ denotes the true class of pixel *i*. The total cross-entropy loss 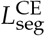 is the mean of 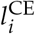 over all pixels and training samples. In results shown in this article, we used equal weights for the two loss terms (i.e. total loss 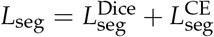). The segmentation model was initialized with a separate copy of the contrastively pretrained encoder weights. Thus, the body segmentation and 3D keypoint models were trained independently but shared the same pretrained initialization. Optimization was performed using the Adam optimizer ^73^. As in the keypoint model, we used a slower learning rate of 3 × 10^−5^ for the pretrained encoder Enc and a faster learning rate of 3 × 10^−4^ for the segmentation decoding head Enc_seg_. The weight decay hyperparameter was set to 10^−5^.

### Inverse/forward kinematics and behavioral replay in NeuroMechFly

We performed inverse kinematics (estimating joint angles from predicted 3D keypoint positions subject to anatomical constraints) and forward kinematics (recomputing anatomically constrained keypoint positions from joint angles) using SeqIKPy ^65^. The anatomical constraints consisted of (i) the available degrees of freedom at each joint and (ii) fixed leg segment lengths. Leg segment lengths were estimated by averaging the distances between corresponding joint keypoints in the raw *PoseForge* predictions over time. To validate the accuracy of the *z* predictions from *PoseForge*, we verified that leg segment lengths in the raw *PoseForge* predictions remained approximately constant over time and were left-right symmetric **(Extended Data Fig. 3)**. The reconstructed 3D pose of the animal with and without anatomical constraints is shown in **Video 4**.

We replayed sequences of inferred joint angles during walking in NeuroMechFly. Only leg joints were articulated. Among these, joints that are actively controlled in the animal (thorax-coxa, coxa-trochanter, femur-tibia, tibia-tarsus) ^30^ were actuated using position actuators to track the recorded motion, whereas compliant joints (between tarsal links) relied on passive joint stiffness and damping. Empirically tuned physics parameters included passive joint damping (0.5 µN mm s rad^−1^), actuator position gain (150 µN mm rad^−1^), sliding friction coefficient (2), and adhesion forces at the leg tips (1 µN for front and middle legs and 0.6 µN for hind legs. These forces were injected at contact points in the normal direction and were divided among multiple contact points when present on the same leg. These parameters were tuned using a Tree-structured Parzen Estimator (TPE) ^76^ implemented in the Optuna hyperparameter optimization library ^77^. During this process, 100 simulations with different physics parameter sets were evaluated to improve the match between simulated and recorded trajectories. These simulations also served as a sensitivity analysis: although certain parameter combinations yielded visibly better trajectory matches, the overall pattern remained robust to moderate variations (within a factor of 4 to 15, depending on the parameter; **Extended Data Fig. 4**). Other untuned parameters are default values from NeuroMechFly (implemented in the FlyGym library, v2.0.0) and MuJoCo ^78^, including passive joint stiffness (10 µN mm rad^−1^), torsional friction coefficient (1), simulation timestep (0.1 ms; recorded kinematics were linearly interpolated from 330 Hz to 10 kHz to match this timestep), and integrator choice (semi-implicit Euler method). Simulations were run on an 8-core Intel® Core™ i9-11900K processor. The physics parameter tuning process took ~ 6 minutes (walltime) with four threads.

The measured trajectory was rotated about the origin to align with the simulated trajectory using the Kabsch algorithm ^79^ to account for different starting orientation. Trajectories shown in **Figure 3b** and **Video 6** were smoothed using a Savitzky-Golay (SG) filter with a 0.5 s window (raw trajectories are also shown in dashed lines in **Figure 3b**). Linear speeds and headings were decomposed from the fly’s velocity in world coordinates, and turning rates were further derived from changes in heading. Linear speeds were computed after applying an SG filter with a 0.5 s window to the raw trajectory, whereas headings and turning rates were computed after applying an SG filter with a 1 s window (a larger window was used for turning rate because it represents a second derivative and is therefore intrinsically noisier).

### Extracting time series of muscle activities

Body segment masks were obtained from *PoseForge* predictions aligned to behavioral camera frames. To refine the mapping between these masks and muscle imaging data, we computed planar homography between the two cameras. Corresponding calibration points were obtained using a custom ChArUco calibration cube imaged simultaneously in both views. The homography matrix *H* mapping behavior-frame coordinates *x*_*b*_ to muscle-frame coordinates *x*_*m*_ was estimated from these correspondences such that *x*_*m*_ ≈ *Hx*_*b*_.

Projected masks were denoised using morphological operations. Fragmented body segment masks containing more than 40 pixels and whose centroids were separated by less than 300 pixels were merged by computing their convex hull. Masks were subsequently dilated to ensure coverage of the underlying muscle regions. For GFP control recordings, masks were expanded using a circular structuring element of radius 7 pixels. For GCaMP recordings, Front leg masks were dilated using custom asymmetric kernels that expanded the masks toward the bottom left and bottom right of the image for the right and left front leg femurs, respectively. With this we avoided expanding toward the bright neck muscles, thereby minimizing contamination of the fluorescence signal. Middle and hind leg masks in the GCaMP recordings were sufficiently precise and were therefore not dilated.

Muscle images were denoised using a bilateral filter (diameter *d* = 5, *σ*_color_ = 150, *σ*_space_ = 150). Within each segment mask, the 500 brightest pixels were selected to form an activity mask. Mean fluorescence was then computed from the raw image intensities of these pixels:

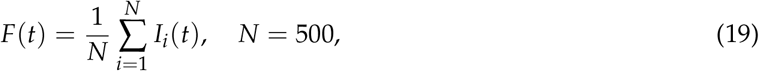

where *I*_*i*_(*t*) is the raw intensity of pixel *i* at time *t*.

Baseline fluorescence *F*_0_ was estimated as the minimum fluorescence value within a 0.5 s window before or after the mechanical stimulation during periods in which the fly was immobile (cumulative stage displacement *<* 0.5 mm). Relative fluorescence changes (signal-over-baseline ratio, Δ*F*/*F*) were then computed as

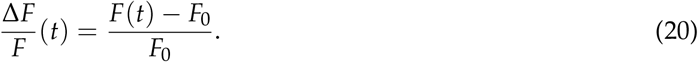

Mechanical stimulation was delivered with a small vibration motor (LilyPad Vibe Board, SparkFun Electronics, USA) and controlled by the Arduino microcontroller that generates all other trigger signals.

For visualization, muscle images were normalized using the 50th–98th percentile range of all recorded pixel intensities during the experiment. The upper bound was further offset by 20 intensity units to suppress visual noise while enhancing contrast in the displayed 16-bit fluorescence images, whose full dynamic range is impractical to render directly. Fluorescence traces over multiple vibration pulses are overlaid relative to vibration onset.

### *Drosophila* husbandry

long-tendon muscles were targeted using the driver line *w[1118]; GMR74F07-GAL4* (Bloomington Drosophila Stock Center, BDSC #39864), kindly provided by Jonathan Enriquez (École normale supérieure de Lyon). Calcium activity was reported using the GCaMP line, *20XUAS-IVS-RSET-jGCaMP8m* (BDSC #605072). The GFP expression control line (genotype *yw; sp/CyO[Dfm::YFP]; 20xUAS-DSCP-6xGFP/TM6C, Tb, Sb*) was kindly provided by Brian McCabe (EPFL).

Experimental GCaMP8m flies (**Figure 4a**) had the genotype *+; +/20XUAS-IVS-RSET-jGCaMP8m; +/GMR74F07-GAL4*. GFP control flies (**Figure 4b**) had the genotype *+; +; 20xUAS-DSCP-6xGFP/GMR74F07-GAL4*.

All flies were reared at 25 ^°^C and 50 % relative humidity under a 12 h light:12 h dark cycle. Experimental animals were recorded 2–5 d post-eclosion.

## Code availability

Hardware design for *Spotlight* is available at https://github.com/NeLy-EPFL/spotlight-hardware. The C++ code base for *Spotlight* real-time control, including embedded Arduino code, is available at https://github.com/NeLy-EPFL/spotlight-control. The Python code base for *Spotlight*-related tools that do not run in real time, including calibration models and postprocessing scripts, are available at https://github.com/NeLy-EPFL/spotlight-tools. *PoseForge* is available at https://github.com/NeLy-EPFL/poseforge. Code used to generate specific figures and videos presented in this paper is available at https://github.com/NeLy-EPFL/spotlight-poseforge-paper. Hardware design for *Spotlight* is licensed under CERN Open Hardware Licence Version 2—Strongly Reciprocal (CERN-OHL-S-2.0). All software are licensed under GNU General Public License Version 3 (GPL-3.0).

## Extended data figures

**Extended Data Fig. 1.**
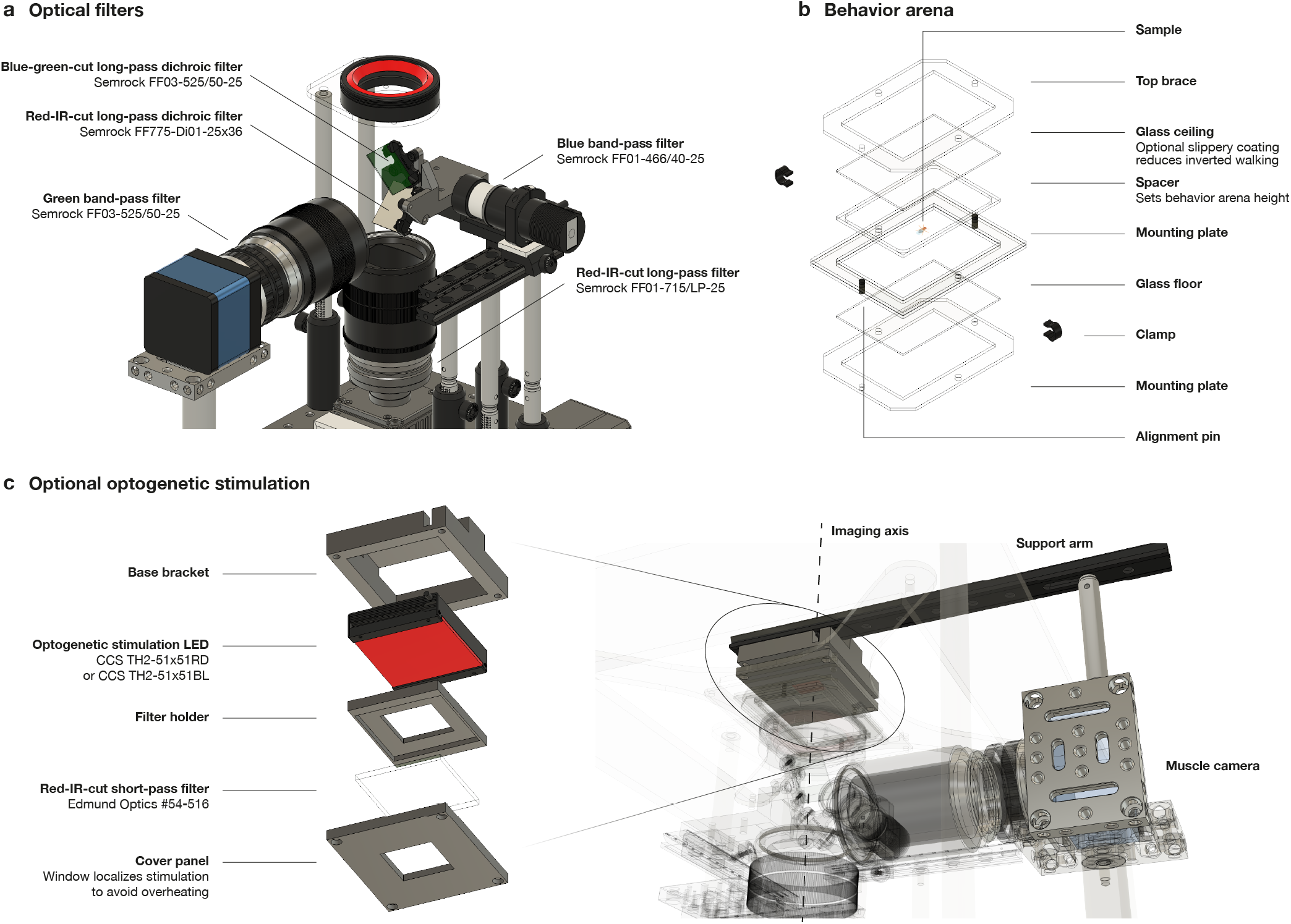
Additional specifications for *Spotlight* construction. **(a)** Placement of optical filters used to separate the light paths for behavior and muscle recording cameras. **(b)** Design and assembly of the behavior arena. **(c)** An additional LED mounted on the muscle camera provides optogenetic stimulation above the arena. See also **Video 2**.

**Extended Data Fig. 2.**
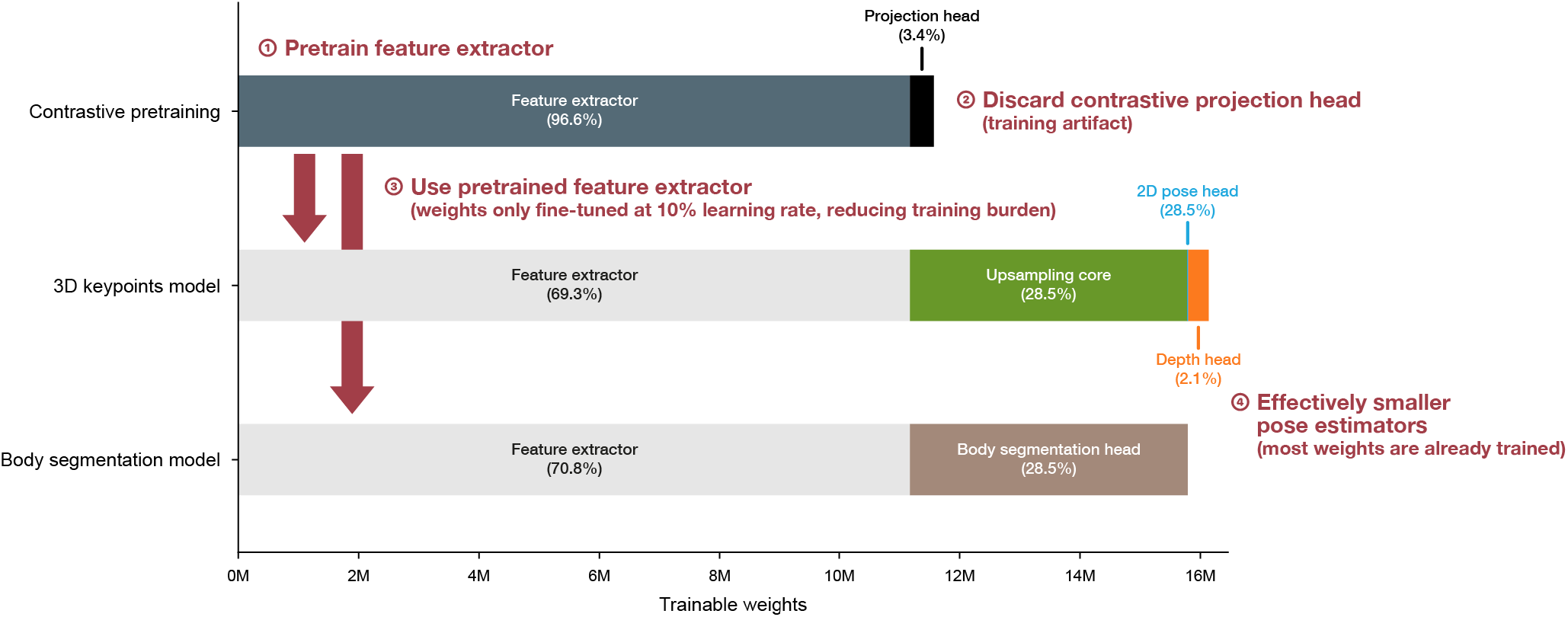
Training procedure and distribution of trainable parameters across model components. X-axis shows the number of neural network parameters optimized during training. Percentages in parentheses indicate the fraction of total trainable parameters from each component. **(Step 1)** During contrastive pretraining, the image encoder (dark blue) is trained through self-supervision. A small projection head (black) maps the general-purpose behavior embedding to an auxiliary latent space where the contrastive loss is applied. **(Step 2)** The contrastive projection head is discarded after pretraining, as it serves only to construct the contrastive loss used to train the encoder. **(Step 3)** The pretrained image encoder then serves as the backbone for the 3D keypoint and body segmentation models, accounting for ~ 70% of their trainable parameters (gray). **(Step 4)** Training can therefore focus on the task-specific heads (green, light blue, orange, brown), while the encoder weights are still fine-tuned with a learning rate equal to 10% of that used for the heads.

**Extended Data Fig. 3.**
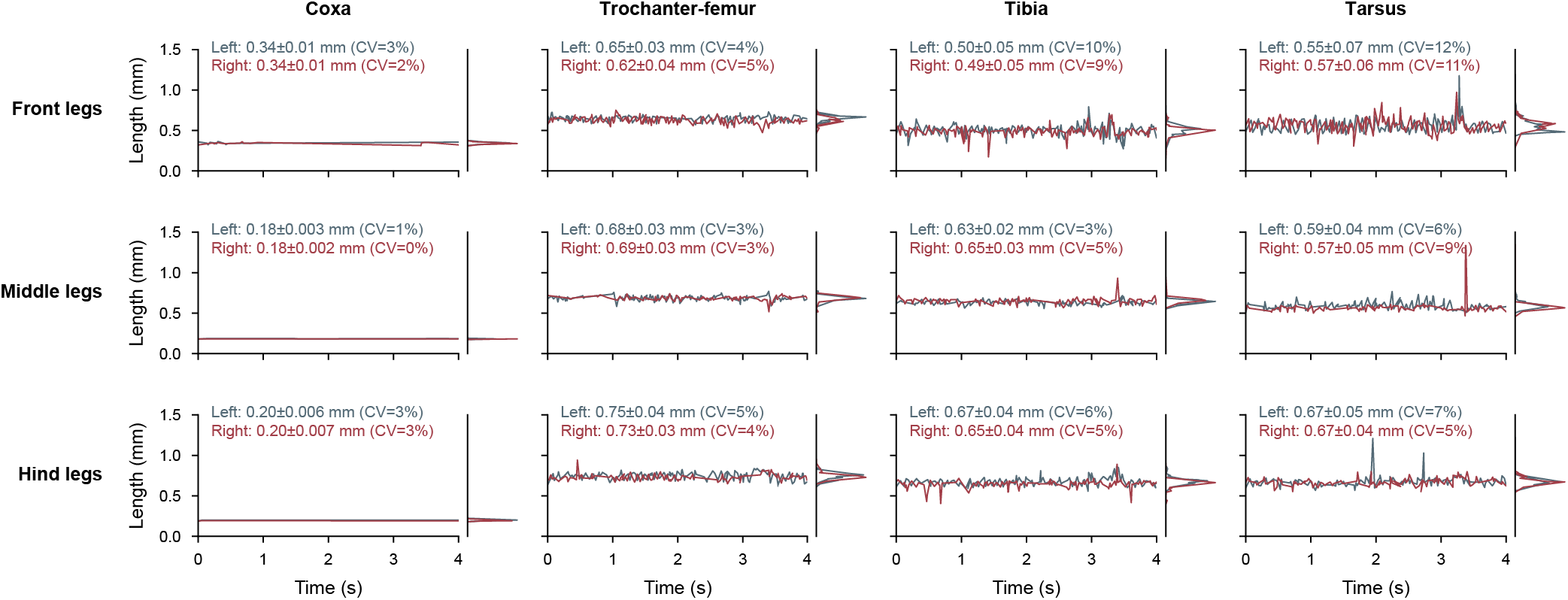
Variation in leg segment lengths in raw *PoseForge* predictions. Larger panels show the temporal variation of leg segment lengths upon median filtering with window size 5 (~ 15 ms), while smaller panels on the right show their distribution. Left and right segments are shown in blue and red, respectively. The time window matches that shown in **Video 4** and **Video 5**. Text annotations indicate the mean and standard deviation of segment lengths, as well as the coefficient of variation (CV; ratio of standard deviation to mean).

**Extended Data Fig. 4.**
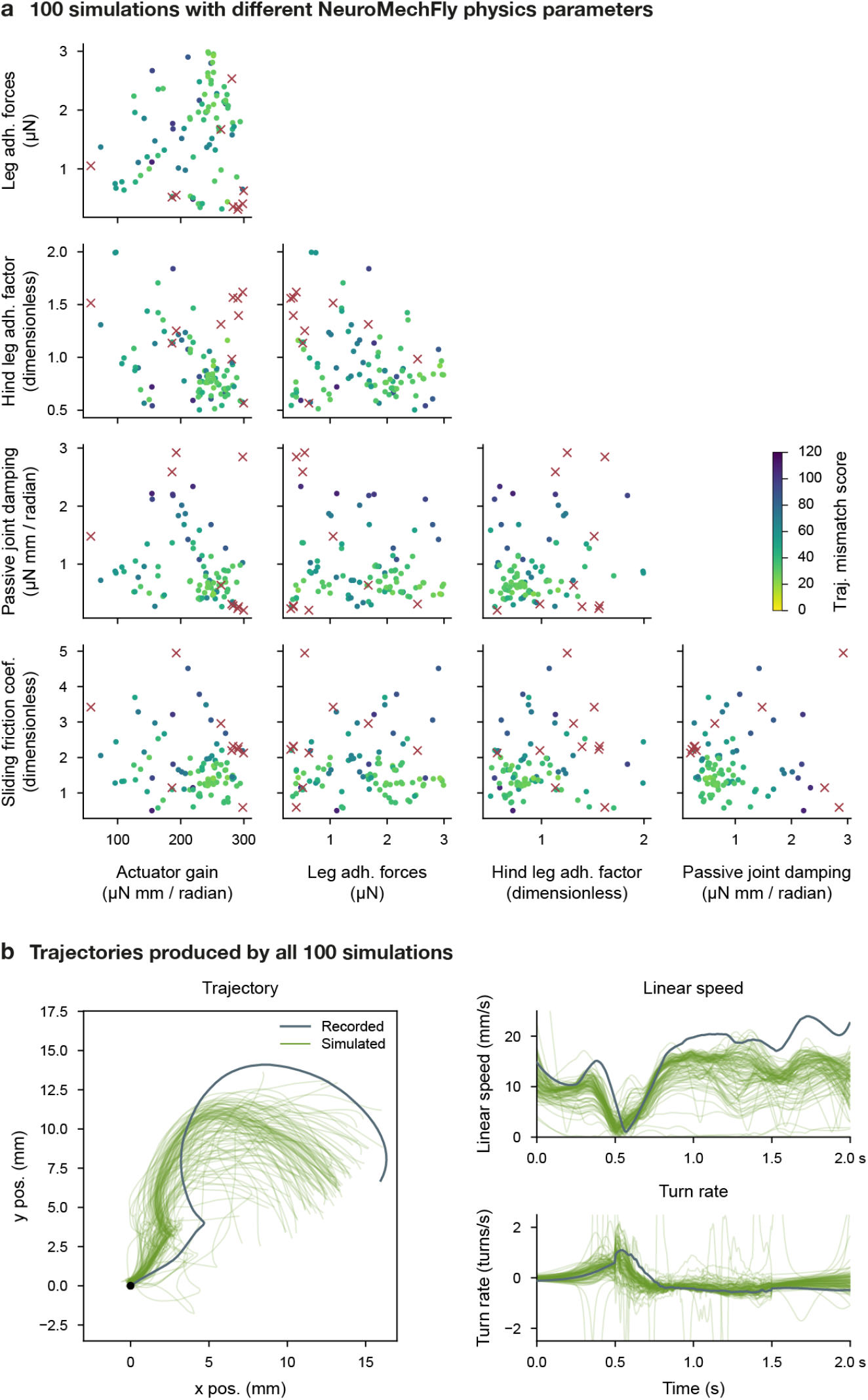
Sensitivity analysis of NeuroMechFly simulations. Physics parameters in NeuroMechFly were varied using a Tree-structured Parzen Estimator (TPE) sampler to explore parameter regimes that improve reconstruction of the recorded fly trajectory. **(a)** Distribution of sampled physics parameters. Marker colors indicate the mismatch between recorded and simulated trajectories (weighted sum of mean squared errors in linear speed and turn rate). Trials with a mismatch score above 150 are considered failed (red crosses). Although clusters of high-performing parameter clusters are visible, the simulated trajectory is generally robust to variations in the varied physics parameters. **(b)** Trajectories generated by simulations (green) compared with the recorded trajectory (blue). Trajectories were rotated to their best alignment to avoid oversensitivity to small heading errors at the beginning of the simulation.

**Extended Data Fig. 5.**
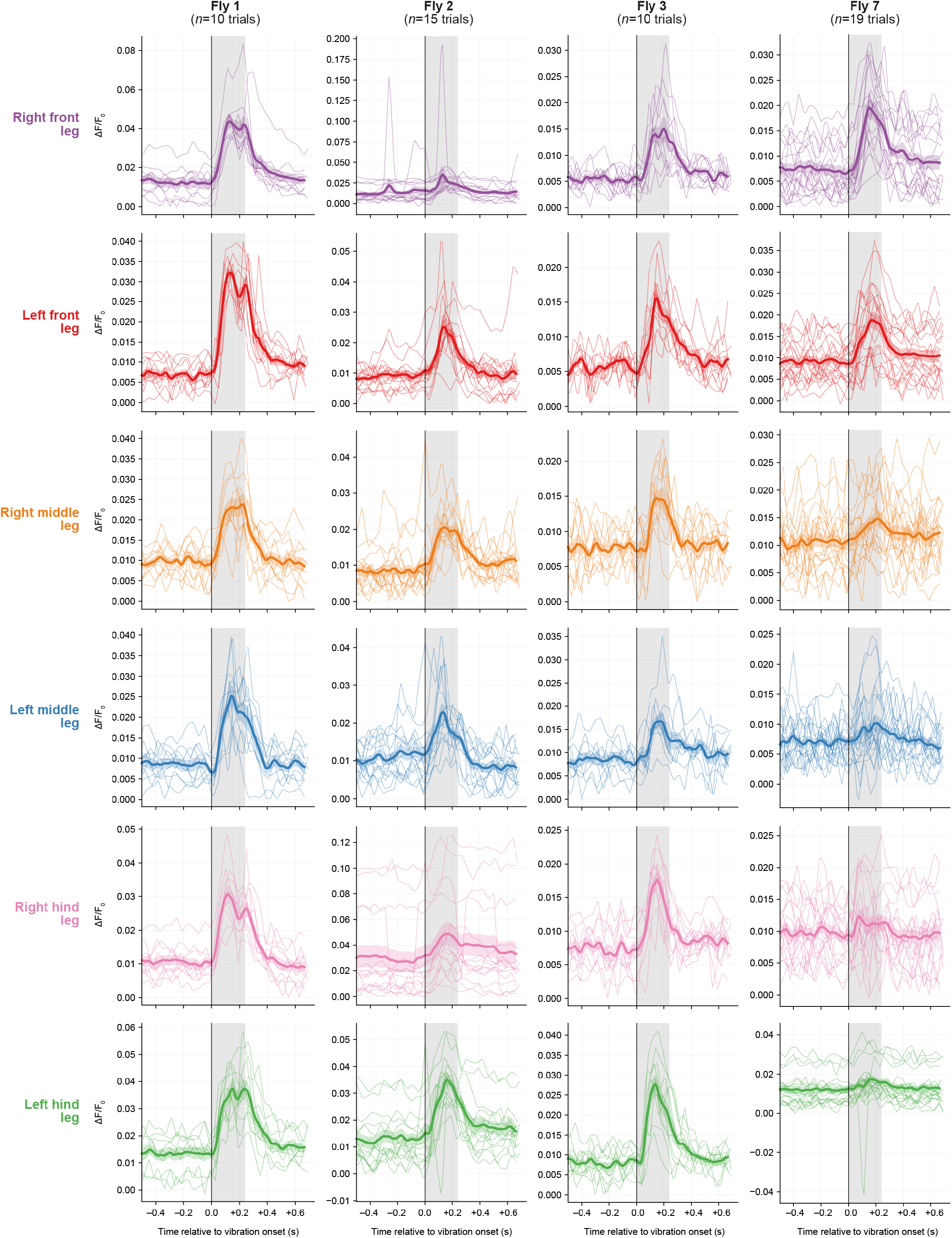
Long-tendon muscle activation pattern over multiple trials. The fluorescence traces of **Figure 4** across multiple vibration pulses are overlaid relative to vibration onset. Vibration-on periods are indicated in gray. Shaded bands indicate mean ± standard error of the mean (SEM).

## Videos

**Video 1: Example video data from the *Spotlight* system**.

Example behavior recording (left) and simultaneous muscle imaging (right) from *Spotlight*.

**Link:** https://go.epfl.ch/spotlight-poseforge#vid1

**Video 2: Design of the *Spotlight* system**.

Overview of the optical and mechanical design of *Spotlight*. See also **Extended Data Fig. 1**.

**Link:** https://go.epfl.ch/spotlight-poseforge#vid2

**Video 3: Example synthetic data and contrastively pretrained latent space for behavior**.

**(top row, left)** Kinematics from a published behavior dataset ^57^ rendered in NeuroMechFly. Ground-truth 2D pose from the simulation is overlaid. **(top row, middle)** Ground-truth 3D pose obtained from the NeuroMechFly simulation. **(top row, right)** Ground-truth segmentation masks corresponding to individual body segments. **(middle rows)** Synthetic videos generated from NeuroMechFly renderings. Panels show variants produced by eight generators with different hyperparameters. **(bottom row)** Latent-space behavior trajectories extracted by an image encoder. Lines with different colors correspond to each of the eight synthetic video variants. The left panel shows trajectories extracted by a generically pretrained encoder; the right panel shows trajectories extracted by the same encoder pretrained on synthetic *Spotlight*-like videos.

**Link:** https://go.epfl.ch/spotlight-poseforge#vid3

**Video 4: Example 3D keypoint positions predicted by *PoseForge***.

Video shown at 0.2× speed; poses are raw predictions without smoothing or filtering. **(left)** Real *Spotlight* recording with 2D pose predicted by *PoseForge* overlaid. **(right)** Raw 3D pose predicted by *PoseForge* (colored lines) and biomechanically constrained 3D pose obtained through inverse and forward kinematics (black lines). Circles indicate the positions of the antennae.

**Link:** https://go.epfl.ch/spotlight-poseforge#vid4

**Video 5: Example body segmentation maps predicted by *PoseForge***.

Video shown at 0.2× speed; poses are raw predictions without smoothing or filtering. **(left)** Real *Spotlight* recording. The apparent shakiness results from per-frame rotation and cropping used to align the fly upright; it can be removed by inverting the alignment transform. **(middle)** Body segmentation maps predicted by *PoseForge*. **(right)** Pixel-level uncertainty estimates produced by *PoseForge*.

**Link:** https://go.epfl.ch/spotlight-poseforge#vid5

**Video 6: Replay of reconstructed behavior in NeuroMechFly**.

**(left)** Real *Spotlight* recording (top) and the experimentally recorded fly trajectory (bottom). **(middle)** The same behavior replayed in NeuroMechFly using position control of the joints. A small delay is expected as forces are generated to match the measured joint angles; the magnitude of this delay depends on the physics (e.g., it may be larger for legs in swing). **(right)** A 3D perspective view of the replayed behavior in NeuroMechFly.

**Link:** https://go.epfl.ch/spotlight-poseforge#vid6

**Video 7: Long-tendon muscle activities upon vibration of the arena in flies expressing GCaMP8m in these muscles**.

Periodic mechanical vibrations were applied using a small vibrator attached near the behavior arena. Top row shows behavior images (left), muscle images (middle), and muscle images with *PoseForge*-extracted masks overlaid (right). Darker pixels on the masks indicate the brightest pixels within the ROI used to extract fluorescence traces. Bottom row shows the Δ*F*/*F*_0_ (signal-over-baseline) fluorescence traces of long-tendon muscles in all legs. Vibration-on periods are indicated in yellow.

**Link:** https://go.epfl.ch/spotlight-poseforge#vid7

**Video 8: Long-tendon muscle fluorescence traces upon vibration in control animals expressing GFP in these muscles**.

Same as **Video 7**, but in GFP control animals where fluorescence levels do not depend on muscle activity levels. Fluorescence remains largely constant during vibration.

**Link:** https://go.epfl.ch/spotlight-poseforge#vid8

## Acknowledgments

SWC acknowledges support from a Boehringer Ingelheim Fonds PhD fellowship. PR acknowledges support from a Swiss National Science Foundation (SNSF) Project Grant (207806). The authors thank Jonathan Enriquez (École normale supérieure de Lyon) for providing the long-tendon muscle driver lines and Brian McCabe (EPFL) for providing the GFP control lines. Fly stocks obtained from the Bloomington Drosophila Stock Center (NIH P40OD018537) were used in this study. Facilities of the EPFL Scientific IT and Application Support Center were used for computation.

## Author Contributions

S.W.-C.—conceptualization, methodology, software, validation, formal analysis, investigation, data duration, writing—original draft, writing—review & editing, visualization.

V.A.S.—conceptualization, methodology, software, validation, formal analysis, investigation, data duration, writing—original draft, writing—review & editing, visualization.

M.A.—methodology, validation, investigation, writing—original draft, writing—review & editing.

P.R.—conceptualization, methodology, resources, writing—original draft, writing—review & editing, supervision, project administration, funding acquisition.

## Ethical compliance

All experiments were computational and thus performed in compliance with relevant national (Switzerland) and institutional (EPFL) ethical regulations.

